# *Flamenco* plasticity tunes somatic piRNAs, rewiring host isoforms and opening a route to heritable transposon spread

**DOI:** 10.1101/2025.10.13.681963

**Authors:** Mai Moritoh, Chikara Takeuchi, Yurika Namba, Hidenori Nishihara, Shigeki Hirakata, Chie Owa, Shinichi Morishita, Yuka W. Iwasaki, Mikiko C. Siomi

**Affiliations:** Department of Biological Sciences, Graduate School of Science, The University of Tokyo, Tokyo 113-0032, Japan; RIKEN Center for Integrative Medical Sciences, Yokohama 230-0045, Japan; Cecil H. and Ida Green Center for Reproductive Biology Sciences, University of Texas Southwestern Medical Center, Dallas, TX 75390-8511, USA; Department of Advanced Bioscience, Faculty of Agriculture, Kindai University, Nara 631-8505, Japan; Department of Computational Biology and Medical Sciences, Graduate School of Frontier Sciences, The University of Tokyo, Chiba 277-8561, Japan

## Abstract

Transposons drive genome innovation, yet how they evade somatic piRNA defenses, reach the germline, and rewire host genes with limited cost remains unclear. Using comparative long-read sequencing in *Drosophila* ovarian somatic cells (OSCs), we show that the LTR transposon *Springer* rewires gene expression by inserting into promoter-proximal introns at an AT-rich motif. Its 5′ LTR initiates transcripts that splice into host exons, expanding isoform diversity without adding coding sequence. We catalog 72 *Springer*-driven isoforms, indicating broad mutagenic potential. In parallel, the *Flamenco* (*Flam*) uni-strand piRNA cluster undergoes structural remodeling that replaces antisense transposon fragments with forward-oriented copies, reshaping piRNA populations and eroding silencing of specific elements, including *Springer*. This relaxation of somatic repression creates conditions permissive for germ-cell entry, providing a plausible route to heritable genome change. Thus, *Flam* plasticity, contrasting with the relative stability of dual-strand clusters, mechanistically links transposon activity to genome rewiring and evolutionary innovation.

## Introduction

Transposons (hereafter referred to as TEs) are pervasive genomic parasites that threaten genome integrity across all domains of life (*1*, *2*). Their mobilization disrupts essential genes and, when occurring in the germline, compromise fertility and transmit deleterious mutations (*3*, *4*). Animals counter this threat with the PIWI-interacting RNA (piRNA) pathway, a small RNA system that enforces TE silencing and preserves reproductive fitness (*5–8*).

piRNAs derive from long single-stranded precursors transcribed from discrete genomic loci known as piRNA clusters (*9–11*). In *Drosophila*, these clusters fall into uni-strand and dual-strand types (*10*, *12*). Both act as genomic archives of fragmented TEs, generating antisense piRNAs that target active elements. Dual-strand clusters employ non-canonical RNA polymerase II (Pol II) and associated cofactors for bidirectional transcription and are restricted to germ cells (*13–15*). Ovarian follicle cells lack this machinery and rely solely on uni-strand clusters, most prominently *Flamenco* (*Flam*) (*12, 16–19*).

Loss of *Flam* function de-represses somatic TEs, establishing *it* as their master suppressor. Because follicle cells encapsulate germ cells, horizontally transmitted TEs first encounter these somatic cells, where *Flam* captures and silences them (*10*, *12*, *20*‒*22*). Cross-species long-read sequencing revealed that *Flam* is variable even among closely related species, reflecting rapid evolutionary diversification (*23*). Some *Drosophila* species lack a syntenic *Flam* yet harbor *Flam*-like loci that repress LTR-type TEs encoding an envelope (Env) protein, paralleling *D. melanogaster* (*24*). These loci are enriched for antisense-oriented LTR-type TE insertion.

Host defenses constrain TE activity, but many TEs persist and can drive regulatory and evolutionary innovation. Realizing such benefits requires germline establishment, yet developmental barriers and robust somatic piRNA defenses sharply restrict access. How, then, do TEs evade piRNA surveillance to colonize the germline? Once established, they can be vertically transmitted across generation (*25*).

Cultured *Drosophila* ovarian somatic cells (OSCs), composed solely of follicle cells (*25*), provide a tractable system to prove these dynamics. We previously discovered that the OSC gene *Lethal* (*3*) *malignant brain tumor* [*L*(*3*)*mbt*] harbors an intronic insertion of the LTR-type TE *Springer*, which drives hybrid splicing (TE-host fusion transcript) from the 5′ LTR to exon 4, generating a truncated isoform, *L*(*3*)*mbt-S* (*26*). Loss of *L*(*3*)*mbt* derepresses piRNA amplification factors and causes female infertility (*26–28*). Yet OSCs tolerate *Springer*, offering an unprecedented window into TE-host interplay.

Here, comparable long-read sequencing reveal that *Springer* reshapes host gene expression through intronic insertions that initiate transcription and drive hybrid splicing without contributing coding sequence. Beyond *L*(*3*)*mbt*, we identified 71 *Springer-*driven isoforms, underscoring its broad mutagenic potential yet modest mutagenicity. *Springer* preferentially integrates into open, promoter-proximal introns, while *Flam* undergoes dynamic remodeling that shuffles TE fragments and reshapes piRNA output and silencing capacity. This plasticity contrasts with the stability of dual-strand clusters in OSCs. It is crucial for protecting both somatic and germline genomes, as TEs with Env can be transmitted from follicle cells to germ cells. In parallel, germline TE integration fuels evolutionary innovation. Thus, *Flam* emerges not only as a fortress against TEs but also as an engine of genome rewiring. Our findings reveal how bursts of TE expression, permitted by *Flam* remodeling, may drive evolutionary novelty.

## Results

### Intronic *Springer* initiates transcription and hybrid splicing without adding coding sequence

To investigate the mechanism underlying OSC-specific *L*(*3*)*mbt-S* isoform (*26*), we generated a haplotype-resolved OSC genome by PacBio HiFi long-read sequencing integrated the data with Micro-C XL profiles (*29*). Contigs were assembled with Hifiasm and scaffolded with RagTag, yielding two haplotypes: one containing the canonical *L*(*3*)*mbt* gene (FlyBase; FBgn0002441) and the other carrying a *Springer* insertion in intron 3 (L3-*Springer*) (Fig. 1A). We hereafter refer to the former as *Haplotype_1* (Hap_1) and the latter as *Haplotype_2* (Hap_2). For convenience, chromosomes other than 3R harboring *L*(*3*)*mbt* were also assigned to Hap_1 or Hap_2.

**Fig. 1.**
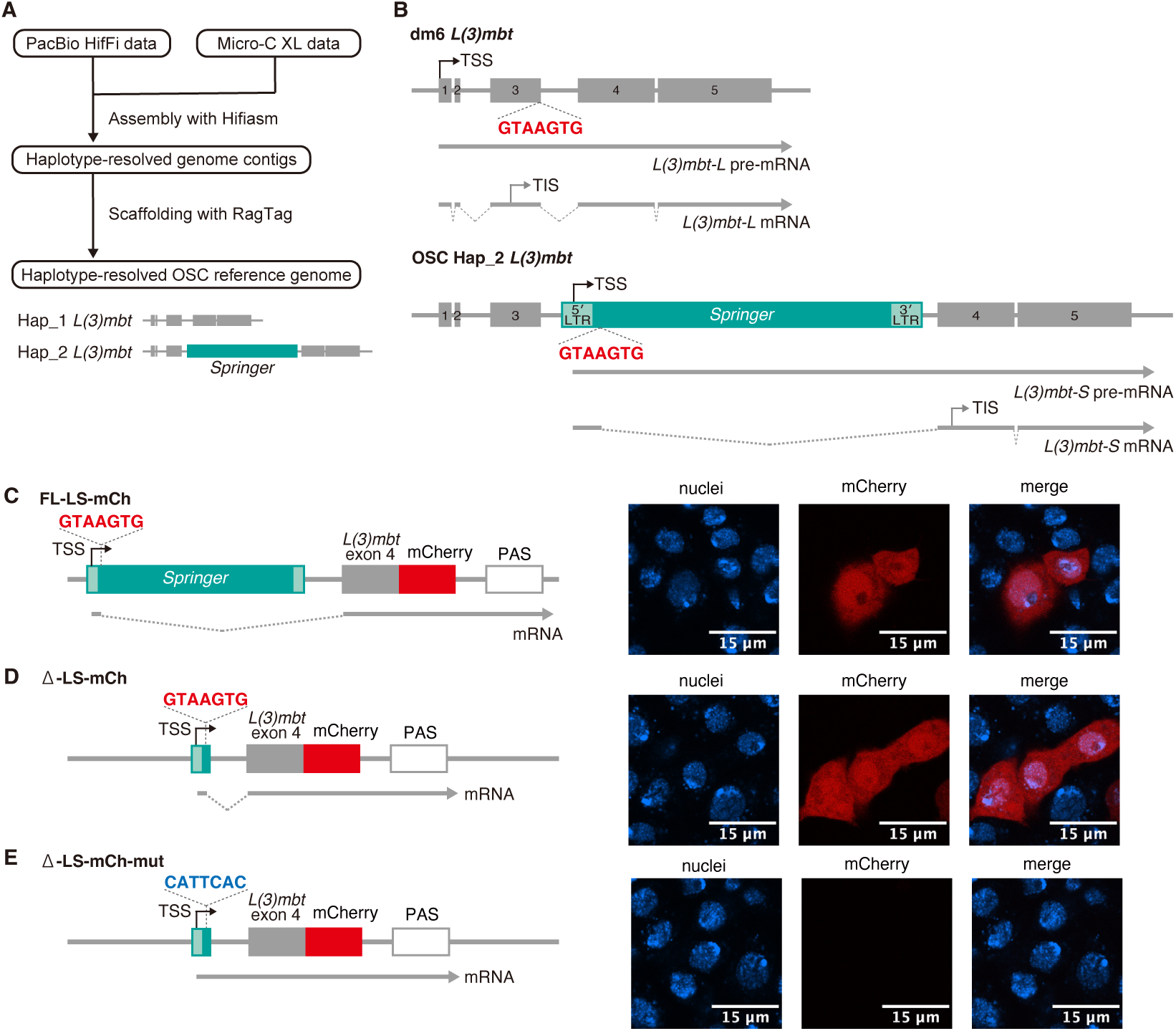
L3-*Springer* introduces a 5′ splice site that drives hybrid splicing to generate the truncated. **L**(**3**)**mbt-S isoform.** (**A**) Genome assemblies of OSC haplotypes: Hap_1 carries intact *L*(*3*)*mbt*, while Hap_2 harbors L3-*Springer* within the locus. (**B**) Structures of *L*(*3*)*mbt* in dm6, Hap_1, and Hap_2. Boxes represent exons (gray) and *Springer* (green); the L3-*Springer* 5′ splice site (GTAAGTG) is highlighted. TSS; transcription start site. TIS; translation initiation site. (**C** to **E**) Plasmid-based recapitulation of *Springer-*driven hybrid splicing: FL-LS-mCh supports splicing and mCherry expression (C); deletion of *Springer* body (D-LS-mCh) preserves expression (D); mutation of the 5′ splice site abolishes expression (D-LS-mCh-mut) (E).

Alignment of Hap_2 with *L*(*3*)*mbt-S* (*26*) revealed a 7-nt splice donor (GTAAGTG) located 43 nt downstream of the 5′ LTR of L3-*Springer* (Fig. 1B). Transcription initiates within the 5′ LTR, omitting exons 1‒3 and including a 297-nt *Springer* fragment that lacks an AUG start codon. Translation therefore begins at Met311 (or Met325) in exon 4, yielding L(3)mbt-S, an N-terminally truncated isoform relative to the full-length L(3)mbt-L (*26*). Thus, L3-*Springer* elicits cryptic transcription initiation and hybrid splicing yet contributes no peptide sequence to the product. Hap_1 expresses L(3)mbt-L as in *Drosophila melanogaster* reference genome (BDGP Release 6 plus ISO1 MT/dm6; GenBank assembly GCA_000001215.4) (Fig. 1B).

Functionally, *L*(*3*)*mbt-S* compensates for loss of *L*(*3*)*mbt-L* in OSCs by repressing targets such as *Vasa* and *Aub*, which are piRNA amplification factors, indicating that N-terminal truncation preserves core activity (fig. S1A). Notably, L(3)mbt-S resembles its vertebrate orthologs (fig. S1B), supporting the hypothesis that TEs can drive evolutionary innovation. Nevertheless, we cannot exclude the possibility that the absence of the N-terminus represents a more primitive state.

*Springer* (FlyBase; FBte0000333; 7,541 nt), an endogenous errantivirus (*30*), produces Gag/Pol from unspliced RNAs and Env from spliced RNAs (fig. S2A). Long-read RNA-seq from OSCs detected both unspliced and spliced *Springer* RNAs and confirmed use of the GUAAGUG donor without auxiliary *Springer* sequences (fig. S2B). Despite an intact polyadenylation signal (PAS) in the 3′ UTR (fig. S2C), polyadenylated L3-*Springer* RNAs were not detected in OSC RNA-seq. In S2 cells, however, L3-*Springer* was efficiently polyadenylated (fig. S2D), indicating that the PAS is intrinsically functional but not recognized in the OSC context. L3-*Springer* harbors a splice acceptor site (GTTACAG) (fig. S2B). Although the core CAG motif is maintained, the upstream 5′ adenine is suboptimal (*31*), likely reducing the splicing efficiency. This may represent an additional mechanism that permits *L*(*3*)*mbt-S* expression in OSCs.

A genomic reporter spanning the L3-*Springer* 5′ end through *L*(*3*)*mbt* exon 4 (FL-LS-mCh) yielded robust mCherry fluorescence and the expected junctions in OSCs (Fig. 1C and fig. S3, A and B). A minimum construct (Δ-LS-mCh) retaining the GTAAGTG donor also supported splicing, whereas mutating this donor (Δ-LS-mCh-mut harboring CATTCAC) abrogated fluorescence (Fig. 1, D and E and fig. S3, A, C, and D), demonstrating that the identified donor is necessary and sufficient for hybrid splicing.

### *Springer*-driven hybrid splicing pervasively rewires host isoforms in OSCs

The dm6 genome contains six full-length or near-full-length *Springer* copies (>7,500 nt; >99.5%) (Fig. 2A and table S1). In OSCs, we identified 219 insertions in Hap_1 and 209 in Hap_2; none matched the six dm6 loci (Fig. 2, B and C). Of these, 150 (Hap_1) and 147 (Hap_2) insertions were genic; 129 (Hap_1) and 126 (Hap_2) were intronic, with 64 and 57 oriented forward relative to host transcription (*Springer*-intF; Fig. 2C and table S2). Analysis of all 428 *Springer* insertions (219 in Hap_1 and 209 in Hap_2) showed a marked enrichment in intronic regions (Fig. 2D), consistent with stronger negative selection against exonic disruption.

**Fig. 2.**
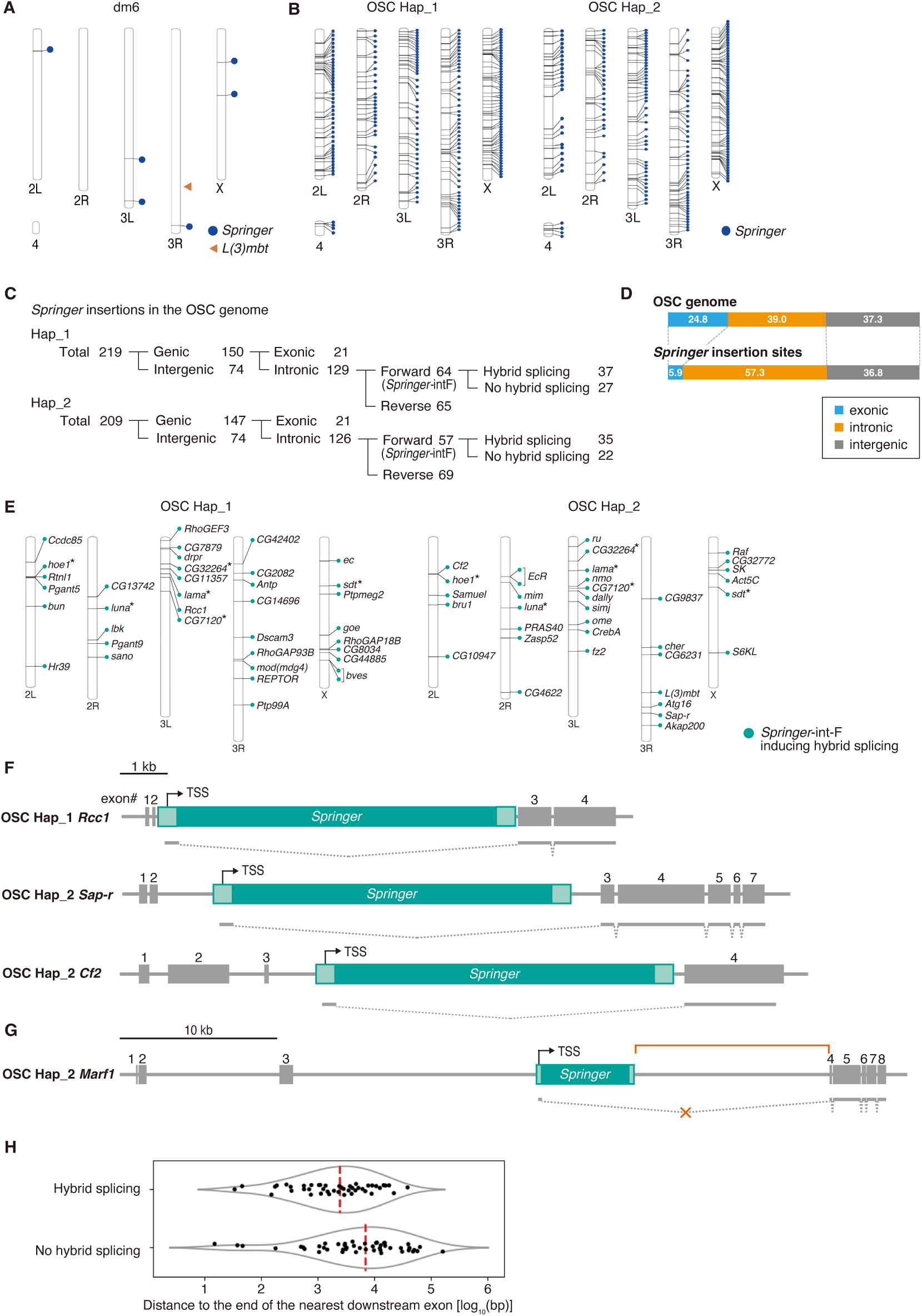
Intronic *Springer* elements broadly promote hybrid splicing and create OSC-specific isoforms. (**A** and **B**) Distribution of *Springer* elements ≥7.5 kb in dm6 and OSC genomes. The *L*(*3*)*mbt* locus in dm6 is also shown. (**C**) Classification of OSC *Springer* insertions. (**D**) *Springer* insertions are enriched in introns. (**E**) Genomic mapping of hybrid splicing *Springer* insertions. *; genes with *Springer* in both haplotypes.]; genes with two *Springer*. (**F**) Examples of host genes (*Rec1, Sap-r, Cf2*) generating *Springer*-induced isoforms. (**G**) *Marf1* carries an intronic *Springer* but shows no fusion transcripts. (**H**) Distances between *Springer* and downstream exons distinguish splicing-competent (Hybrid splicing) versus non-competent insertions (No hybrid splicing). Red bar; means. Mann–Whitney U test, p = 5.022e−05.

Long-read RNA-seq revealed hybrid splicing for 37 of 64 (Hap_1) and 35 of 57 (Hap_2) *Springer*-intF sites, including L3-*Springer* (Fig. 2, C, E, and F, fig. S4A, and table S3). Five genes carried *Springer*-intF at identical positions in both haplotypes, implying uniform expression of N-terminally truncated isoforms in OSCs (Fig. 2E). In contrast, dm6 carries no *Springer*-IntF (fig. S4B) and lacks short *Springer* segments akin to Δ-LS-mCh (Fig. 1D), suggesting that *Springer*-dependent isoforms are specific to OSC genomes.

### Long downstream introns disfavor hybrid splicing at *Springer*-intF

Roughly 30% of *Springer*-intF insertions [27 (Hap_1) and 22 (Hap_2)/121] showed no hybrid splicing (Fig. 2, C and G, fig. S4C, and table S4). These sites ley farther from the nearest downstream exon: median 9,572 nt without splicing vs 2,661 nt with splicing (D = 6,911 nt; Fig. 2H). We propose competition between splicing and premature polyadenylation: long downstream introns favor PAS usage within *Springer* before the acceptor is reached, yielding *Springer* RNAs rather than hybrid isoforms. Additional parameters, Pol II elongation kinetics and acceptor strength, likely modulate this balance.

### OSCs produce abundant but sense-biased *Springer* piRNAs

Although ovarian *Springer-*piRNAs effectively silence *Springer* (*32*, *33*), OSCs harbor numerous *Springer* insertions. Reanalysis of OSC piRNAs (*34*) showed that *Springer-*piRNAs are abundant (Fig. 3A), yet Piwi depletion failed to derepress *Springer* (fig. S5A). Interestingly, *Springer* piRNAs in OSCs are strongly sense-biased (Fig. 3, B and C), consistent with limited silencing efficacy. *Copia*, *gypsy1*, *Transpac*, and *Xanthias* showed similar sense biases and, like *Springer*, were not markedly derepressed in Piwi-deficient OSCs (Fig. 3, B and C and fig. S5A), indicating that residual antisense piRNAs play little role in their silencing. This is consistent with a previous report showing that antisense piRNAs in limited amounts are insufficient to trigger silencing (*35*).

**Fig. 3.**
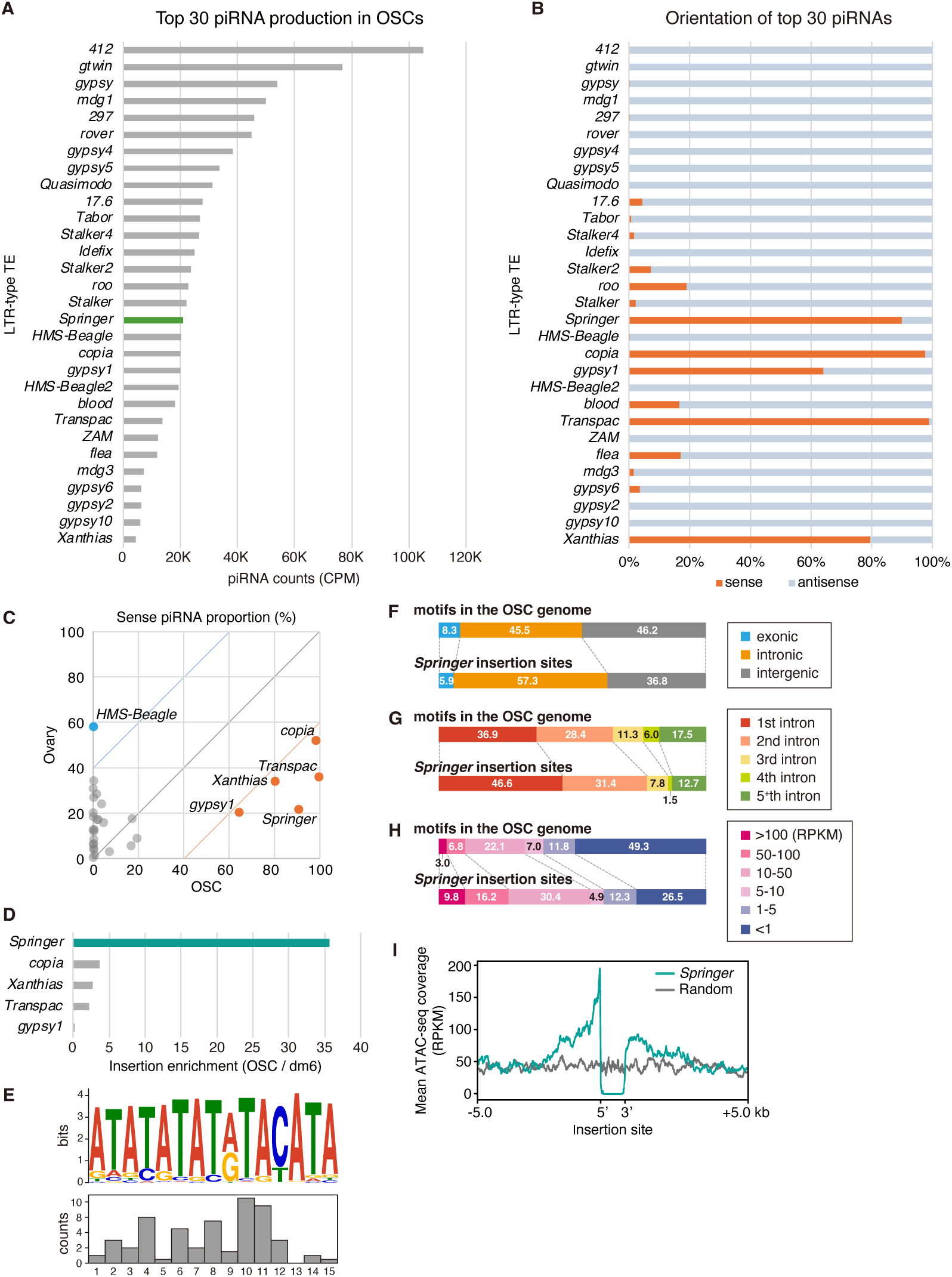
OSCs are rich in sense-oriented *Springer-*piRNAs and allow accumulation of *Springer* elements in active, promoter-proximal introns. (**A** and **B**) Piwi-bound piRNAs mapped to LTR TEs in OSCs, highlighting sense bias for *Springer*. (**C**) Comparative analysis with ovaries shows *Springer, copia, Transpac,* and *Xanthias* acquire sense piRNAs in OSCs, unlike most other TEs. (**D**) *Springer* exhibits the strongest genomic enrichment, followed by *copia, Xanthias,* and *Transpac*. (**E**) Sequence motif enriched at *Springer* insertion sites. Bar graph shows insertion frequency within the motif. (**F** and **G**) *Springer* insertions are strongly biased toward introns, particularly near 5′ ends of genes. (**H**) Insertions preferentially occur in highly expressed regions. (**I**) ATAC-seq analysis indicates *Springer* preference for open chromatin. Random; control.

### *Copia* rarely drives hybrid splicing and often perturbs coding frames

*Springer* showed the highest enrichment, with an ∼36-fold increase in copy number (Fig. 3D). *Copia* increased ∼3-fold, while *Xanthias* and *Transpac* each increased ∼2.5-fold. In contrast, *gypsy1* showed no increase (<1), likely due to abundant *gypsy1*-targeting piRNAs in OSCs (Fig. 3B).

We identified 119 (Hap_1) and 129 (Hap_2) *copia* insertions, with 35 and 45 intronic forward-oriented sites (*copia*-intF; fig. S5B). Only 1/35 (Hap_1) and 4/45 (Hap_2) yielded detectable hybrid splicing (fig. S5C), far below *Springer*-intF frequencies. We attribute this to weaker internal donor strength (GUAUGUU), competition with premature polyadenylation, and potential negative selection. Indeed, *copia*-derived exonic segments are long (1,484 nt; FlyBase; FBte0000023) and contain multiple AUGs, predisposing to frameshifts or fusion peptides that may compromise cell viability. *Xanthias*-intF and *Transpac*-intF showed no hybrid splicing (fig. S5, D and E), consistent with the absence of intrinsic splicing in these elements. Genic *gypsy1* insertions were not detected (fig. S5F). Together, *Springer* uniquely and efficiently drives coding-sequence-free hybrid splicing.

### *Springer* targets open, promoter-proximal introns of active genes

We asked what genomic contexts favor *de novo Springer* integration. We found that the insertions (219 Hap_1; 209 Hap_2) preferentially occurred within a distinctive AT-rich repeat array and showed a central positional bias within this array (Fig. 3E). Genome-wide insertions were enriched in introns relative to exons or intergenic space (Fig. 3F). Although 23.8% of the OSC genome is exonic (Fig. 2D), only 5.9% of *Springer* insertion sites occurred in exons, less than expected from motif frequency, consistent with purifying selection against exon disruption.

Insertions were enriched in first/second introns and in highly expressed genes (Fig. 3, G and H). ATAC-seq meta-profiles centered on insertion sites revealed a sharp accessibility peak immediately upstream of integrations, absent from matched random controls (Fig. 3I), indicating a preference for open chromatin near 5′ gene regions. We hypothesize that transcription initiation and local Pol II density promote *Springer* integration, consistent with known roles of host factors in TE targeting.

### Extensive *Flam* remodeling reshapes piRNA populations and silencing capacity

We compared *Flam* for *Springer, copia, Xanthias, Transpac,* and *gypsy1* fragments between dm6 and OSC haplotypes, focusing on inserts >300 nt due to repetitiveness. In dm6, two reverse-oriented *Springer* fragments (dm6-Sp#1 and #2) produce *Springer*-targeting piRNAs (Fig. 4A and fig. S6A). In OSCs, these are absent; instead, multiple near-full-length, forward-oriented *Springer* fragments populate *Flam* in both haplotypes (Sp#3, Sp#4, Sp#5 in Hap_1 and Sp#4 in Hap_2; Fig. 4, B and C), generating predominantly sense *Springer* piRNAs (fig. S6A) and explaining ineffective *Springer* silencing.

**Fig. 4.**
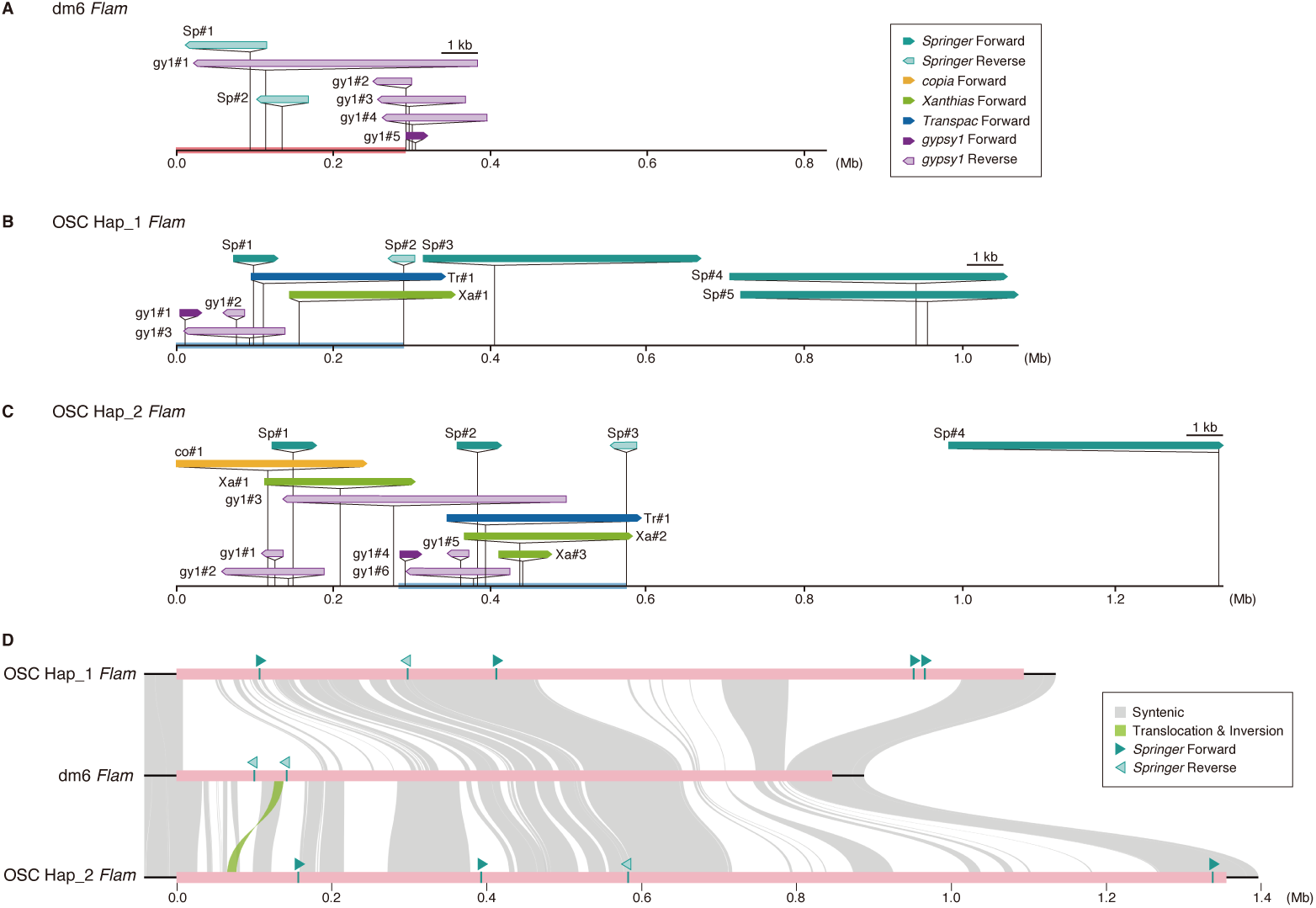
*Flam* emerges as an unparalleled hotspot of genome rearrangement. (**A** to **C**) Comparison of TE fragments in the *Flam* locus among dm6, OSC Hap_1, and Hap_2, with *Springer, copia, Xanthias, Transpac*, and *gypsy1* indicated. (**D**) Structural rearrangements in the *Flam* region and its 50-kb flanking sequences among the dm6 genome and the OSC Hap_1 and Hap_2 assemblies.

*Gypsy1* fragments also revealed remodeling: multiple reverse-oriented copies in dm6 correspond to a reconfigured set across Hap_1/Hap_2, with evidence of a ∼300-kb gap relative to dm6 (Fig. 4, A to C and fig. S6B). *Copia, Xanthias,* and *Transpac* fragments are undetectable in dm6 *Flam* (down to 20-nt searches) but present in OSC *Flam* (Tr#1 and Xa#1 in Hap_1 and co#1, Tr#1, and Xa#1/#2 in Hap_2), which produces corresponding piRNAs (Fig. 3, B and C and fig. S7, A to C). Ovarian piRNAs from these TEs map mainly to dual-strand clusters (table S5). Collectively, *Flam* undergoes pervasive remodeling that rewires piRNA populations and impacts TE activity in OSCs.

### *Flam* is a hotspot of genome rearrangement

Synteny mapping of dm6 against Hap_1 and Hap_2 revealed intact continuity at the boundaries of *Flam* but fragmented synteny across most of the locus (Fig. 4D). Non-syntenic sequence comprised 52.9% (vs Hap_1) and 38.9% (vs Hap_2) of dm6 *Flam*, including a minor translocation/inversion event. Although Hap_1 and Hap_2 are broadly similar, Hap_2 uniquely harbors a ∼330-kb 5′ segment densely populated with TE fragments (Fig. 4D and fig. S8A). Dot-plots further showed that, while *Flam* in dm6 and OSC (Hap_1 or Hap_2) is rich in structural variation, the remainder of the X chromosome is comparatively quiescent (Fig. 5, A to F): 93.5% and 92.6% of the dm6 non-*Flam* X retain synteny with Hap_1 and Hap_2, respectively, even across typically unconstrained noncoding regions.

**Fig. 5.**
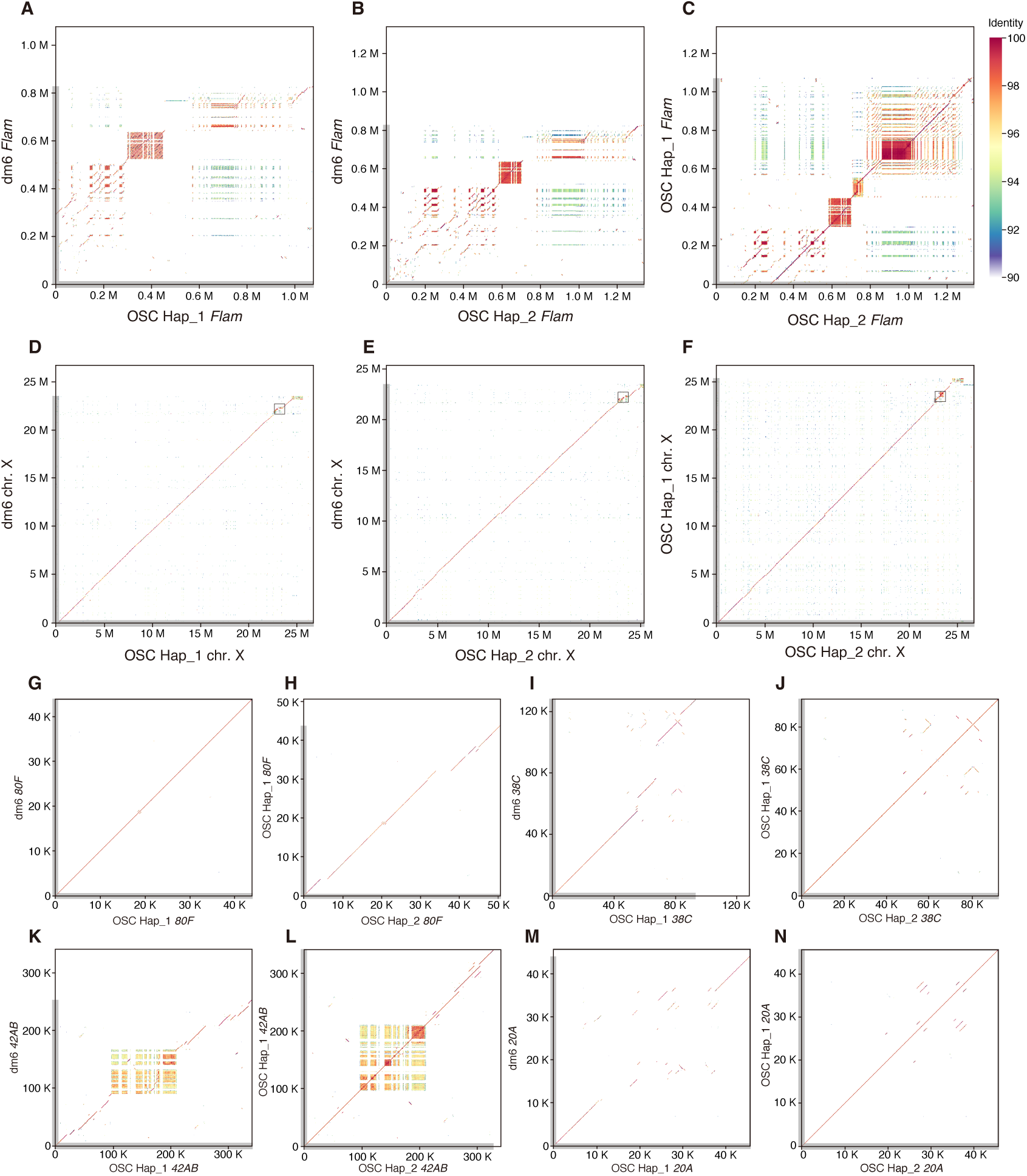
The *Flam* locus is rearrangement-prone compared with other genomic regions. (**A** to **C**) Pairwise comparisons of *Flam* between dm6, Hap_1, and Hap_2 assemblies reveal large-scale rearrangements. (**D** to **F**) The X chromosome shows limited rearrangements relative to *Flam*, underscoring *Flam*’s exceptional plasticity. The areas indicated by open boxes correspond to the data in A to C. (**G** and **H**) Pairwise comparisons of dual-strand cluster *80F*. (I and J) Pairwise comparisons of *38C*. (**K** and **L**) Pairwise comparisons of *42AB*. (**M** and **N**) Pairwise comparisons of *20A*.

Among major dual-strand clusters [*80F* on 3L (∼44kb, dm6), *38C* on 2L (∼127kb)*, 42AB* on 2R (∼253kb)] (*10, 13*) (fig. S8B), *80F* is largely invariant, whereas *38C* and *42AB* show modest variation (Fig. 5, G to L and fig. S8, C to E). Chromatin accessibility is relatively low at 38C but higher at *80F* and *42AB* compared with their flanking regions (fig. S9A to C), showing a partial correlation with rearrangement frequency that appears largely independent of clusters usage in OSC piRNA biogenesis.

The tiny dual-strand cluster *20A* (∼44kb, dm6), proximal to *Flam*, is more variant than *80F* and *38C* (Fig. 5, M and N and fig. S8F), and more open (fig. S9D), yet the intervening ∼120-kb *20A*-*Flam* interval is much less open than either *20A* or *Flam*. Notably, *80F*, *38C*, *42AB*, and *20A* retain complete synteny between Hap_1 and Hap_2, whereas *Flam* exhibits marked haplotype disparity (fig. S8B), establishing *Flam* as a hotspot of genome rearrangement in OSCs.

Comparison of our OSC *Flam* assemblies with prior OSC datasets (*32*) shows uniform read coverage across *Flam* in both haplotypes (Fig. 6A) and only minor differences in *Springer* copy number over the past decade (Fig. 6B) (*32*, *36*). Together, these observations support a model in which *Flam* explains via recurrent local remodeling, rather than through recent large-scale gains or losses.

**Fig. 6.**
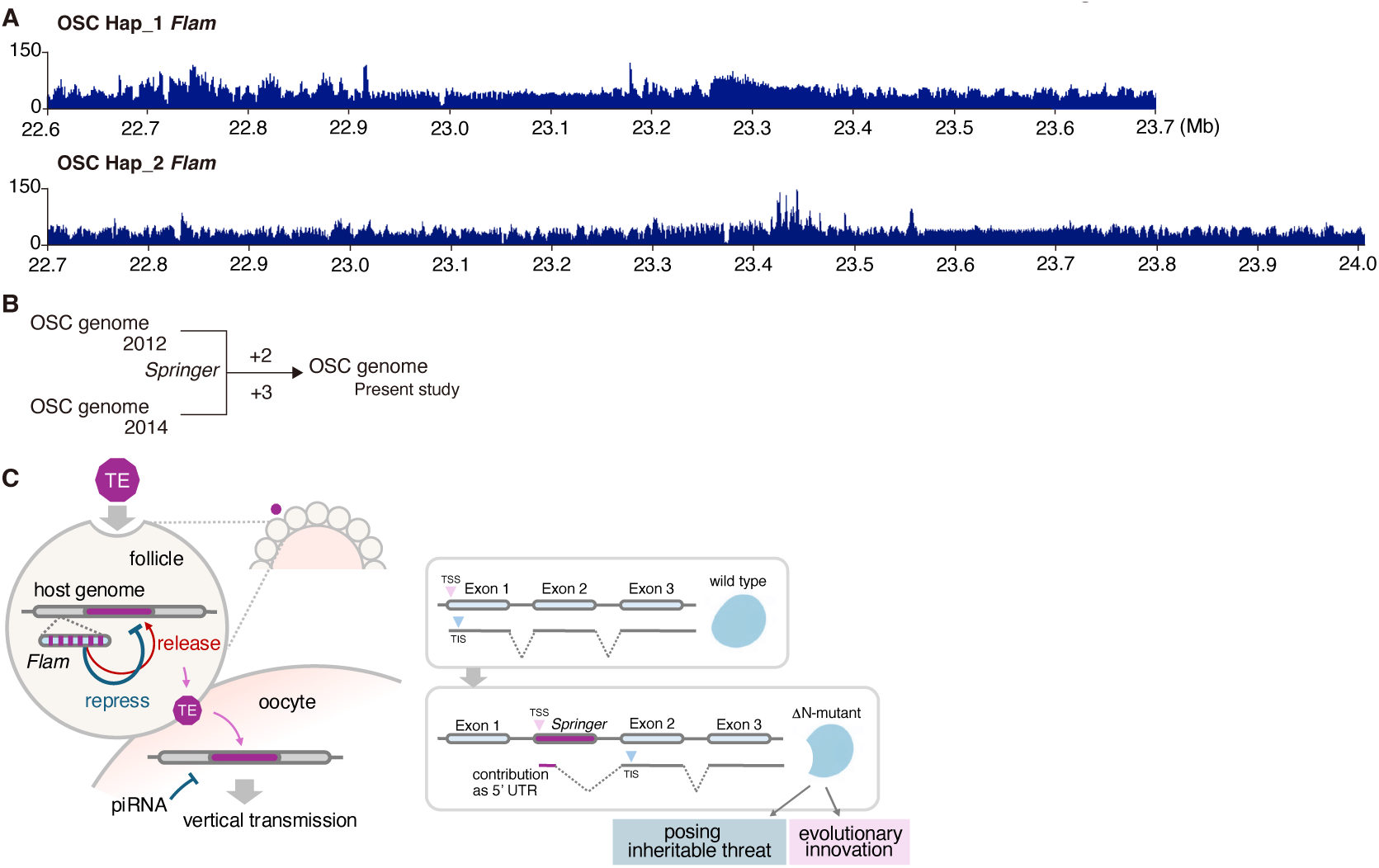
*Flam* robustness in contrast to its plasticity. (A) Comparison of our OSC *Flam* with previously published OSC datasets (*34*). (B) *Springer* insertion sites relative to previous genome assemblies (*34, 35*). (**C**) Model illustrating how *Flam* plasticity couples *de novo* TE insertions to genome rewiring and evolutionary innovation.

## Discussion

Long-read, haplotype-resolved genomics of *Drosophila* OSCs uncover two linked properties that connect TE activity to host gene regulation: a promoter-proximal, splice-competent strategy of LTR element *Springer*, and a *Flam* piRNA cluster that is unusually prone to local remodeling. Together, these features provide a mechanistic route by which somatic defenses can be retuned and host isoforms rewired̶an often-invoked but rarely resolved path to genome innovation. At the same time, our coverage and copy-number analyses indicate no large-scale gain or loss at *Flam* over decadal sampling, suggesting recurrent local rearrangements rather than wholesale turnover.

*Springer* preferentially integrates into AT-rich, promoter-proximal introns. Its 5′ LTR acts as a cryptic promoter whose transcripts splice into host exons without adding coding sequence, thereby expanding isoform diversity while imposing minimal proteomic cost. In contrast to elements whose exonic contributions introduce ATGs and fusion peptides̶features likely subject to purifying selection̶*Springer*’s strategy decouples transcriptome diversification from proteome expansion. Within our assays and detection limits, no other tested elements showed comparable, coding-sequence-free hybrid splicing at similar frequency, positioning *Springer* as a prototypical driver of this mode of regulatory rewiring.

Although intronic TE‒host hybrids from *de novo* TE insertions have been reported, these were isolated cases most detected with short reads (*37*‒*41*). Here, we provide the first systematic dissection of a *de novo* TE insertion and its functional consequences in one cell type, uncovering an unrecognized mechanism that positions *Springer* as a genome-wide driver of regulatory innovation. Experimental evidence underscores this potential: insertion of a *Springer* mini-cassette (Δ-LS-mCh, Fig. 1D) into *Piwi* intron 2 abolished its nuclear localization signal encoded across exons 1 and 2, converting Piwi into a cytoplasmic protein (fig. S10). Such alterations may ultimately feed into evolutionary change.

*Flam*, by contrast, replaces reverse-oriented fragments with forward-oriented copies, reshaping piRNA populations toward sense bias and weakening element-specific silencing in OSCs. Because derepressed TEs in follicle cells can, in principle, access neighboring germ cells, such somatic relaxation may create conditions permissive for germline entry and eventual heritable change (Fig. 6C). We emphasize that this is a mechanistic inference supported by directionality and piRNA data; direct measurement of germline integration rates and multigenerational transmission will be important future tests.

Open questions also include the molecular basis of *Flam*’s remodeling, the host factors guiding *Springer*’s promoter-proximal preference, and the extent to which similar coding-sequence-free hybrid splicing operates across elements, strains, and species. Resolving these will clarify how genomes balance defense with innovation̶not by turning TE activity off, but by shaping when and how it is allowed to matter.

## Materials and Methods

### Cell culture

OSCs (*25*) (DGRC Stock 288; https://dgrc.bio.indiana.edu//stock/288; RRID:CVCL_IY73) were cultured at 26°C in Shields and Sang M3 Insect Medium w/ L-Glutamine (US Biological) supplemented with 10% fly extract (*25*), 10% fetal bovine serum (FBS; NICHIREI), 10 mU/mL insulin, and 0.6 mg/mL glutathione. S2 cells were cultured at 26°C in Schneider’s *Drosophila* Medium (Gibco) with 10% FBS.

### Genome sequencing

For HiFi sequencing, genomic DNA was extracted from OSC using a Genomic-tip 20/G (QIAGEN) and a Genomic DNA Buffer Set (QIAGEN). To assess the quality and integrity of the genomic DNA samples, we ran them for sequencing on Pippin Pulse pulsed-field electrophoresis (Sage Science). We used 0.75% agarose gel (SeaKem Gold, Lonza) on a pre-set default program of 5‒80kb (75 V, 16 h run time) with approximately 500 ng DNA/lane. According to the PN101-853-100 protocol (PacBio), the genomic DNA was sheared using a Megaruptor 2 (Diagenode), and the fragments were collected using 0.45x AMPure PB beads (PacBio). The size of the DNA fragments was checked using a TapeStation System (Agilent Technologies). SMRTbell libraries were prepared using a SMRTbell Express Template Prep Kit 2.0 (PacBio) and a BluePippin DNA Size Selection System (Sage Science). The selected libraries were bound with Sequencing Primer v2 and Sequel II Polymerase 2.0 using a Sequel II Binding Kit 2.0 (PacBio). Sequencing was performed on a Sequel II instrument (PacBio) in CCS mode using a SMRT Cell 8 M and a Sequencing Plate 2.0. This process included eight hours of pre-extension and 30 hours of movie viewing.

### Assembly of the OSC genome

A haplotype-resolved genome of the OSC cell line was assembled using the Hifiasm software (*42*). This assembly was performed utilizing PacBio long-read sequencing data generated in this study, in conjunction with publicly available Micro-C XL data from OSC cells (*29*). The resulting contigs were subsequently aligned and scaffolded against dm6 reference genome with the Ragtag toolkit (*43*). For all subsequent analyses, the OSC genome assembly was defined as the primary contigs mapping to chromosomes 2L, 2R, 3L, 3R, 4, and X were used, unless otherwise specified.

### OSC gene annotation

To annotate the OSC genome, gene annotations were transferred from the dm6 reference genome. First, we generated chain files to map coordinates between the dm6 reference and each OSC haplotype (Hap_1 and Hap_2) using the flo workflow (*44*). Using these chain files, we then transferred the dm6 gene annotations (UCSC) to our Hap_1 and Hap_2 assemblies with the liftOver utility from the UCSC toolkit (*45*).

### RNAi and transfection

For RNAi, OSCs were first electroporated with 200 pmol of siRNA in 20 μ L of Solution SF (Cell Line Nucleofector Kit SF, Lonza) using a Nucleofector 96-well Shuttle device (Lonza). Following electroporation, cells were left at room temperature for 10 min before plating into culture dishes, and then incubated at 26 °C for 48 h. Cells were subsequently subjected to a second electroporation with 600 pmol of siRNA dissolved in 100 μ L of transfection buffer (180 mM sodium phosphate, pH 7.2, 5 mM KCl, 15 mM MgCl₂, and 50 mM D-mannitol) using the Nucleofector 2b system (Lonza). After a 10-min rest at room temperature, the cells were plated again and incubated at 26 °C for an additional 48 h before collection. siRNA sequences are listed in table S7.

For plasmid transfection, 1.0 × 10^7^ OSCs or S2 cells were mixed with 6‒ 15 μ g of plasmid DNA in 100 μ L of the same transfection buffer modified from a previous report (*46*) and electroporated with a Nucleofector 2b device (Lonza). Cells were maintained at 26 °C and harvested 48 h after transfection.

### Nuclear fractionation

Nuclear fractionation was carried out following a modified version of a previously described protocol (*47*). Briefly, cells were suspended in hypotonic buffer [10 mM HEPES-KOH (pH 7.3), 10 mM KCl, 1.5 mM MgCl_2_, 0.5 mM dithiothreitol (DTT), 2 μ g/mL leupeptin, 2 μ g/mL pepstatin A, and 0.5% aprotinin], gently mixed by pipetting, and disrupted by repeated passage through a 25-gauge needle. Lysates were centrifuged at 400 g for 10 min, and the pellet was collected as the crude nuclear fraction, while the supernatant was retained as the cytoplasmic fraction. The cytoplasmic fraction was adjusted to 200 mM KCl and clarified by centrifugation at 20,000 g for 20 min. The nuclear pellet was washed with hypotonic buffer, resuspended in chromatin co-immunoprecipitation buffer [50 mM HEPES-KOH (pH 7.3), 200 mM KCl, 1 mM EDTA, 1% Triton X-100, 0.1% sodium deoxycholate, 2 μ g/mL leupeptin, 2 μ g/mL pepstatin A, and 0.5% aprotinin], then sonicated and centrifuged at 20,000 g for 20 min. The resulting supernatant was used as the nuclear extract for western blotting.

### Western blotting

Proteins from nuclear extracts were resolved by SDS‒polyacrylamide gel electrophoresis and transferred onto Immobilon-P membranes (Millipore). Membranes were blocked in PBS containing 0.1% Tween-20 (T-PBS) supplemented with 5% skim milk and then incubated with anti-L(3)mbt antibody (1:1,000 dilution) (*26*). After washing with T-PBS, membranes were incubated with horseradish peroxidase‒conjugated anti-mouse IgG secondary antibody (1:10,000 dilution; Cappel, cat. no. 55558). Protein bands were visualized using Clarity Western ECL Substrate (Bio-Rad), and chemiluminescence signals were captured with a ChemiDoc XRS Plus imaging system (Bio-Rad).

### RT-qPCR

Total RNA was extracted from OSCs using ISOGEN II (NIPPON GENE) and treated with TURBO DNase at a final concentration of 0.04 U/ μ L (Thermo Fisher Scientific) to remove residual genomic DNA. Reverse transcription was performed with ReverTra Ace (Toyobo) according to the manufacturer’s protocol. Quantitative PCR was carried out on a StepOnePlus Real-Time PCR System (Thermo Fisher Scientific) using THUNDERBIRD Next SYBR qPCR Mix (Toyobo). Relative transcript abundance was determined using the ΔΔCt method, with *rp49* as the internal reference gene. Primer sequences are listed in table S7.

### Iso-seq and bioinformatic analysis

Total RNA was extracted from cultured cells using ISOGEN II (NIPPON GENE), followed by DNase treatment, phenol/chloroform purification, and ethanol precipitation. Full-length cDNA libraries were prepared with the SMRTbell Prep Kit 3.0 (Pacific Biosciences) according to the manufacturer’s protocol. Sequencing was performed on a PacBio Sequel II platform in HiFi read mode (1-cell run), generating 4,504,293 HiFi reads comprising 9.73 Gb of sequence data, with a read N50 of 2,234 bp, an average read length of 2,160 bp, and an average read quality of Q45. HiFi reads were processed using the Iso-Seq3 pipeline (v.4.0.0) (https://github.com/ylipacbio/IsoSeq3). Demultiplexing and primer removal were conducted with lima in Iso-Seq mode, specifying custom 5′ and 3′ primer sequences (5′ -GCAATGAAGTCGCAGGGTTGGG-3′ and 3′ -GTACTCTGCGTTGATACCACTGCTT-5′). Full-length non-concatemer (FLNC) reads were refined with the --require-polya option to confirm poly(A) tails, and high-quality consensus isoforms were obtained by clustering with quality value scoring enabled. The curated isoforms were aligned independently to OSC genome Hap_1 and Hap_2 using minimap2 (v.2.24-r1122) (*48*) in splice-aware mode with quality value support, while suppressing secondary alignments. Alignment files were processed with SAMtools (v.1.16.1) (*49*) to convert, sort, and index BAM files for downstream analysis.

### Plasmid construction

For the L3-*Springer* plasmid (fig. S2D), the L3-*Springer* sequence from the OSC Hap_2 genome was PCR-amplified from OSC genomic DNA and inserted into a modified pAc-based vector lacking the promoter region using Mighty Cloning (Takara).

For FL-LS-mChe (Fig. 1C), a genomic fragment spanning the *L*(*3*)*mbt* locus from the L(*3*)-*Springer* insertion site in intron 3 to the putative start codon of exon 4 (OSC Hap_2 genome) was PCR-amplified from OSC genomic DNA. This fragment, together with an mCherry sequence PCR-amplified from pIB_DDX4N_mCherry_CRY2PHR, was assembled into a modified pAc-based expression vector lacking the promoter using NEBuilder HiFi DNA Assembly Master Mix (New England Biolabs). The Δ-LS-mChe (Fig. 1D) and Δ-LS-mChe-mut (Fig. 1E) constructs were generated by inverse PCR.

For the Piwi plasmid (fig. S10), the *Piwi* genomic region (including introns) was PCR-amplified from OSC genomic DNA and assembled into the same modified vector with a C-terminal FLAG tag using NEBuilder HiFi DNA Assembly Master Mix (New England Biolabs). To generate the Piwi+Δ-LS construct (fig. S10), the Δ-LS fragment was amplified from the Δ-LS-mChe template and incorporated into the Piwi plasmid using NEBuilder HiFi DNA Assembly Master Mix. Primer sequences used in these plasmid constructions are listed in table S7.

### 3′ RACE

S2 cells were transfected with the L3-*Springer* plasmid by electroporation using the Nucleofector 2b device (Lonza) with program N-20 and cultured at 26°C for 2 days. Total RNA was extracted with ISOGEN II (NIPPON GENE), followed by DNase treatment and phenol/chloroform extraction with ethanol precipitation. 3′ RACE-ready cDNA was synthesized using the SMARTer RACE 5′ /3′ Kit (Takara Bio) according to the manufacturer’s instructions. The resulting 20 µL cDNA was diluted with 90 µL Tricine-EDTA buffer. PCR amplification was carried out with SeqAmp DNA Polymerase (Takara Bio) using a gene-specific primer containing a 15-bp overlap sequence for cloning (5′ - gattacgccaagcttGCTCCATCTTCAGTGAGCTCGTGTGCGC-3′) under touchdown PCR conditions (Program 1). PCR products were resolved on a 1% TBE agarose gel, and the target band was excised and purified with the FastGene Gel/PCR Extraction Kit (Nippon Genetics). Purified fragments were assembled into the pRACE vector using NEBuilder HiFi DNA Assembly Master Mix (New England Biolabs) and transformed into NEB 5-alpha Competent *E. coli* (New England Biolabs). Colony PCR was performed with SapphireAmp Fast PCR Master Mix (Takara Bio) using the gene-specific primer and M13 forward primer. PCR products were treated with Exonuclease I (New England Biolabs) and Shrimp Alkaline Phosphatase (New England Biolabs) and sequenced using the BigDye Terminator v3.1 Cycle Sequencing Kit (Thermo Fisher Scientific).

### RT-PCR

Total RNA was extracted from OSCs using ISOGEN II (NIPPON GENE) and treated with TURBO DNase at a final concentration of 0.04 U/ μ L (Thermo Fisher Scientific) to eliminate residual genomic DNA. Reverse transcription was performed with the Transcriptor First Strand cDNA Synthesis Kit (Roche) using oligo(dT) primers. The resulting cDNA was amplified with Q5 High-Fidelity DNA Polymerase (New England Biolabs). PCR products were resolved on a 1% TBE agarose gel, stained with SYBR Gold (Thermo Fisher Scientific) for 10 min, and visualized with a transilluminator (fig. S3A). Primer sequences are listed in table S7.

### Live-cell imaging

OSCs were cultured on glass-bottom dishes (MATSUNAMI). Hoechst 33342 (Lonza) was added to the medium at a final concentration of 1 µg/mL, and cells were incubated at 26°C for 30 min. Live-cell imaging was performed as previously described (*50*), using a confocal laser scanning microscope (LSM 980, Carl Zeiss) equipped with a Plan-Apochromat 63×/1.4 Oil DIC M27 objective lens at room temperature. Hoechst 33342 and mCherry were excited with 405 nm and 561 nm lasers, respectively, and images were acquired under identical acquisition settings across all samples within each experiment.

### *Springer* insertion analysis

The dm6 and OSC genomes were analyzed with RepeatMasker (v.4.1.5) (*51*) to annotate TEs. From the resulting GFF files, entries corresponding to the LTR retrotransposon *Springer* were extracted. Insertions ≥7500 nt in length (≥99.5% of the reference consensus length) were classified as full-length *Springer* elements. Genomic insertion sites of these elements were visualized using Phenogram (https://visualization.ritchielab.org/phenograms/plot; Fig. 2, A to C) (*52*). Phenogram was also used to visualize selected insertions in Fig. 2E.

### *Springer* insertion site analysis

We categorized novel *Springer* insertions based on their location relative to the lifted-over gene annotations. Insertions were classified as “genic” if they fell within an annotated exon or intron, and “intergenic” otherwise. Genic insertions were further subdivided into “exonic” or “intronic”. To analyze the distance to the nearest splice site, we focused on intronic insertions located on the sense strand relative to the host gene and calculated the distance from the insertion site to the start of the nearest downstream exon using a custom script. Insertions that were found within transcripts identified by our Iso-seq data, potentially creating TE-gene fusion transcripts, were classified as “hybrid splicing” events (see code availability for details).

For the analysis in Fig. 3, E-H, we annotated the genomic features of both the FIMO-identified motif sites and the novel *Springer* insertion sites by mapping their coordinates to the dm6 reference genome. Feature distribution was determined using annotatePeaks.pl. 3′ UTR, 5′ UTR, exon, and non-coding were categorized as “Exonic”, whereas intergenic, TTS, promoter-TSS were grouped as “Intergenic”.

To assess the expression level of genes targeted by insertions or containing the motif, we used OSC gene expression data from FlyBase (release FB2025_01). The intron number (e.g., first intron, second intron) for each intronic insertion was also determined from the HOMER output (*53*).

### TE insertion site and motif analyses

To identify *Springer* and *copia* insertion sites in the dm6 genome, 100 bp flanking sequences of these insertions from the OSC genome were mapped back to the dm6 reference. The center points of successfully mapped flanking pairs were defined as the insertion site. Annotations of these insertion sites are analyzed using Homer (annotatePeaks.pl) (*53*). *De novo* motif discovery on these insertion sites was performed using MEME-ChIP (*54*). *Springer* motif sites were defined by FIMO (*55*) at the threshold of p=0.001 and annotated in the same way.

### Small RNA analysis

Previously published small RNA-seq datasets were reanalyzed: ovary small RNAs from the *Drosophila melanogaster* strain *y[1]; cn[1] bw[1] sp[1]* (SRR827770) (*56*) and Piwi immunoprecipitation (Piwi-IP) smRNA-seq from OSCs (SRR9158321) (*34*).

For the ovary dataset, the Illumina 3′ adapter sequences (TGGAATTCTCGGGTGCCAAGGAACTCCAGTCAC) was removed using Cutadapt (v.4.6) (*57*). Reads outside the 20‒35 nt size range were discarded, leaving ∼13 million reads for analysis. For the OSC dataset, preprocessing was performed following previous study (*34*), in which four random nucleotides from each end and adapter sequences were removed using Cutadapt, and only reads between 20‒35 nt size range were retained for analysis, resulting in ∼3.7 million reads.

Trimmed ovary reads were aligned to dm6, while OSC reads were mapped to a merged OSC genome assembly (Hap_1 and Hap_2). In both cases, alignments were performed using Bowtie (v.1.3.1) (*58*) with identical parameters (-v 1, ≤1 mismatch; --best), ensuring consistency between datasets. Only reads that successfully mapped to the dm6 or merged OSC genome were retained for subsequent analyses.

For piRNA production analysis (Fig. 3, A to C), genome-mapped reads were further aligned to consensus sequences of *Drosophila* transposable elements obtained from FlyBase using Bowtie with no mismatches allowed, and random multi-mapping was performed with the -k 1 option. Read counts were normalized to counts per million mapped reads (CPM). To quantify strand-specific mapping, custom Python scripts were used to count the number of sense and antisense reads for each transposon, based on SAM alignment flags.

For transposable element (TE) locus‒specific analyses (Figs. S6 and S7), the previously generated alignment files were converted to BAM format, sorted, and indexed using SAMtools. Normalized coverage tracks were then produced with deepTools (bamCoverage, v3.5.5) (*59*) using counts per million mapped reads (CPM) normalization. The resulting BigWig files were visualized in Integrative Genomics Viewer (IGV) (*60*).

### TE enrichment analysis

The dm6and the merged OSC genome assembly (Hap_1 + Hap_2) were annotated with RepeatMasker. From the resulting GFF files, insertions ≥99.5% of the TE consensus length were defined as full-length. For each TE, genomic intervals were merged, the occupied bases were summed, and genome occupancy was calculated as the fraction of occupied bases relative to total genome size. Enrichment was defined as the ratio of occupancy in OSC genome to that in dm6. Custom Python scripts (Biopython-based) were used for these calculations (Fig. 3D).

### ATAC-seq analysis

Previously published ATAC-seq data (*61*) were re-analyzed for this study.

First, raw reads were processed for quality control: read errors were corrected using Rcorrector (v1.0.4) (*62*) and TranscriptomeAssemblyTools (https://github.com/harvardinformatics/TranscriptomeAssemblyTools), and adapter sequences were removed using TrimGalore (v0.6.6) (https://github.com/FelixKrueger/TrimGalore). Reads shorter than 50 bp were subsequently discarded. The filtered reads were aligned to each of the Hap_1 and Hap_2 haplotypes of the OSC genome using Bowtie2 (v2.5.4) (*63*) with the following parameters: “--maxins 4000 --very-sensitive --no-mixed --no-discordant”. Uniquely mapped reads were selected using SAMtools (v1.21). Read coverage was then calculated and normalized to reads per kilobase per million mapped reads (RPKM) using the bamCoverage tool from the deepTools suite (v 3.5.4), setting the bin size to 10 bp and the minimum mapping quality to 10. For datasets with biological replicates, the signal was averaged using the bigwigAverage tool from deepTools to generate a single representative BigWig file per haplotype. These files were used for visualization in IGV (fig. S9). To analyze chromatin accessibility around *Springer* insertion sites (Hap_1: 219 sites; Hap_2: 209 sites), signal distributions were computed using the computeMatrix tool from deepTools in scale-regions mode over 10 kb window (5 kb upstream and downstream of each site), with a 10 bp bin size, treating missing data as zero. As a control, 200 random genomic regions of 7,546 bp (matching the full-length *Springer* element length) were selected using the BEDTools random command (v2.31.1) (*64*) and processed with the same pipeline. Mean signal prorfiles for both *Springer* and control sites were generate mean profile plots with the plotProfile tool from deepTools. The signal profiles for Hap_1 and Hap_2 were then averaged and plotted using R (*65*) (Fig. 3I).

### Definition of piRNA cluster regions

The *Flam* locus was defined as the genomic region between *CG32820* and *CG14621*, which are conserved flanking genes identified in both the dm6 annotation and the lifted-over OSC gene annotations. Other piRNA cluster regions were defined based on the *D. melanogaster* piRNA cluster annotations from the ProTRAC database (*66*), using sequence coordinates corresponding to the reported cluster boundaries. See table S7 for locus information.

### *Flam* structural rearrangement analysis

Structural differences among the dm6 and OSC genome assemblies were analyzed using minimap2 (*48*) and SyRI (*67*). Genome alignments were performed with minimap2 (v2.29) using the “-cx asm5 ‒eqx” option in two runs: OSC Hap_2 assembly was aligned to dm6, and dm6 was aligned to the OSC Hap_1 reference assembly. The resulting PAF files were analyzed with SyRI (v1.7.1) using the “--cigar ‒nosnp” option. Rearrangements in the flam region and its 50-kb flanking sequences were visualized by excluding highly diverged regions, using plotsr (v1.1.1) (*68*) with the “̶ itx” mode.

### Dot plot analysis

Dot plots were generated with ModDotPlot (*69*) to compare, for each genomic region, the following pairs: dm6 vs. OSC Hap_1, dm6 vs. OSC Hap_2, and OSC Hap_1 vs. OSC Hap_2. Unless noted otherwise, default settings were used with an identity threshold of -id 90. The sliding window parameter -r was set per analysis as follows: 600 for the *Flam* locus (Fig. 5, A to C), 1,000 for the X chromosomes (Fig. 5, D to F), 800 for 80F (Fig. 5, G and H and fig. S8C), and 600 for 38C (Fig. 5, I and J and fig. S8D), and 600 for 42AB (Fig. 5, K and L and fig. S8E), and 800 for 20A (Fig. 5, M and N and fig. S8F).

### Genome comparison analysis

DNA-Seq data from previous study (*32*) were mapped to the haploid genome (Hap_1/Hap_2) using Bowtie2 (*63*) with default settings. A TE insertion was considered pre-existing if reads spanning the TE and its flanking sequences were found in the data. Conversely, TE insertions that lacked support from such spanning reads were identified as novel insertions.

### Identification of insertion sites in other OSC/OSS genomes

The genomic DNA-Seq datasets from previous studies (*32*, *36*) are mapped to Hap_1 and Hap_2 respectively using Bowtie2. Then, if the reads which intersect TE and flanking sequencings are detected, these TEs are considered as existing in the genome of the other dataset.

### Immunofluorescence

OSC cells were fixed and processed for immunostaining as described previously (*70*). Cells were incubated with an anti-FLAG monoclonal antibody (M2, Sigma-Aldrich; 1:1,000 dilution) as the primary antibody, followed by incubation with an Alexa Fluor 488‒conjugated anti-mouse IgG secondary antibody (Thermo Fisher Scientific; 1:500 dilution). After washing, samples were mounted in VECTASHIELD Mounting Medium with DAPI (Vector Laboratories) and imaged using a confocal laser scanning microscope (LSM 980, Carl Zeiss).

### Statistics and reproducibility

RT-qPCR (fig. S1A and S5A) was performed over three independent experiments, and other live-cell imaging (Fig. 1, C to E), western blotting (fig. S1A), 3′ RACE (fig. S2D), RT-PCR (fig. S3A), and immunofluorescence (fig. S10) were performed over two independent experiments. Statistical procedures are described in the figure legends. Sample sizes were not predetermined, and experiments were performed without randomization or blinding.

## Acknowledgments

We thank H. Yamazaki, S. Yamanaka, R. Saito, K. Saito, and other members of the Siomi laboratory at the University of Tokyo for useful discussions. We thank I. Oka and T. Morita for technical assistances.

## Funding

Japan Society for the Promotion of Science 24K18093 (YN)

Japan Society for the Promotion of Science 25H01308 (HN)

Japan Society for the Promotion of Science 24K09376 (SH)

Japan Society for the Promotion of Science 25H01305 (YWI)

Japan Society for the Promotion of Science 25H01303 (MCS)

Japan Society for the Promotion of Science Research Fellowship for Young Scientists (DC2) 22KJ2696 (CT)

Fusion Oriented REsearch for disruptive Science and Technology JPMJFR224L (YWI)

## Author contributions

Conceptualization: SM, YWI, and MCS

Methodology: MM, YN, and

Investigation: MM, CT, YN, HN, SH, and CO

Supervision: SM, YWI, and MCS

Writing-original draft: Everybody

Writing-review & editing: YN, YWI, and MCS

## Data and materials availability

The genome assemblies generated in this study have been deposited in the NCBI GenBank under accession numbers XXXX–XXXX. The RNA-seq datasets have been deposited in the NCBI Sequence Read Archive (SRA) under accession numbers SRRXXXXXXX–SRRXXXXXXX. Code and data are available at GitHub (https://github.com/Chikara-Takeuchi/2025_moritoh_TE/tree/master).

## Figures and Tables

**Fig. S1.**
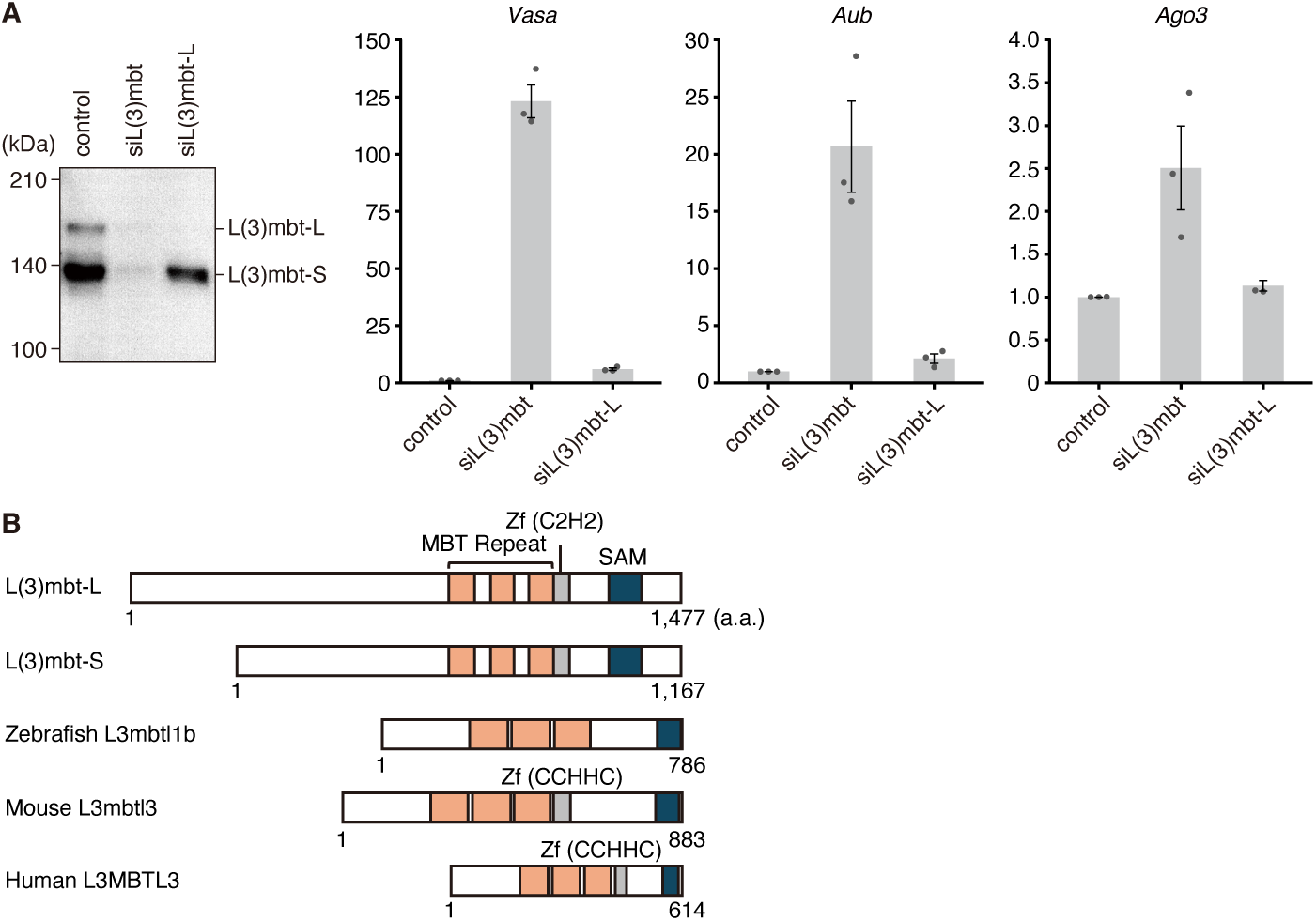
L(3)mbt isoforms in OSCs and conservation across species. (A) Western blot of L(3)mbt-L and L(3)mbt-S in control OSCs, cells depleted of both isoforms [siL(3)mbt], and cells selectively depleted of L(3)mbt-L [siL(3)mbt-L] (left). RT-qPCR (right) shows that L(3)mbt-S retains the ability to repress piRNA amplification factors (*Vasa*, *Aub*, and *Ago3*). (B) Conservation of L(3)mbt homologs: zebrafish L3mbtl1b (Gene ID: 559572), mouse L3mbtl3 (Gene ID: 237339), and human L3MBTL3 (Gene ID: 84456).

**Fig. S2.**
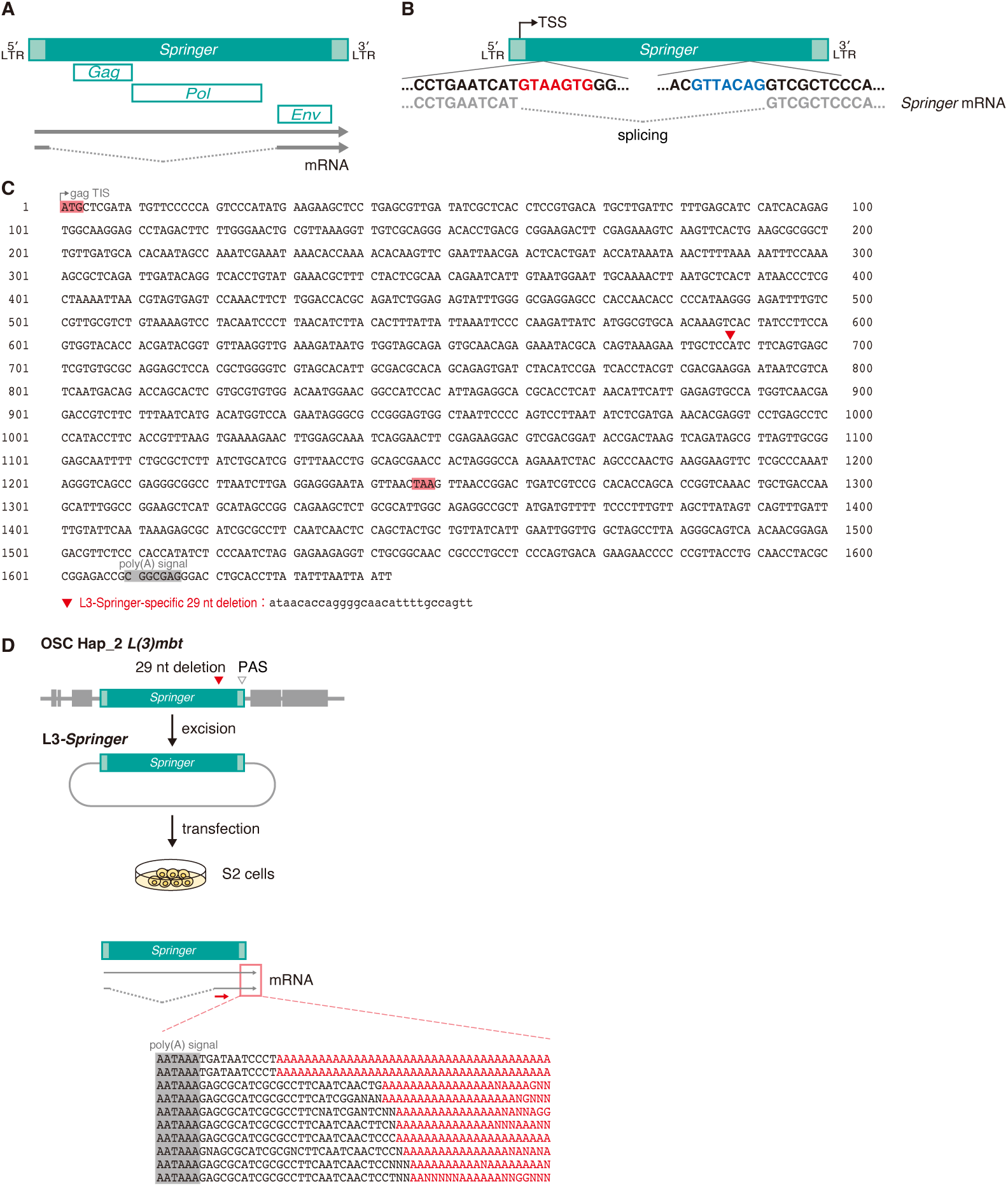
The *Springer* element and L3-*Springer*. (**A**) Schematic of the *Springer* element showing the Gag, Pol, and Env ORFs and the architectures of its unspliced and spliced transcripts. (**B**) The splice donor (red) and splice acceptor (blue) sites within *Springer*. (**C**) DNA sequence of the L3-*Springer* segment from the 5′ end of the *Env* coding region to the 3′ end of L3-*Springer*. The initiator methionine (iMet) and stop codon are highlighted in red; the L3-*Springer*-specific 29-nt deletion is marked with a red triangle. The sequence is shown in lowercase. The poly(A) signal is shaded gray. (**D**) L3-*Springer* expression in S2 cells; transcripts are polyadenylated.

**Fig. S3.**
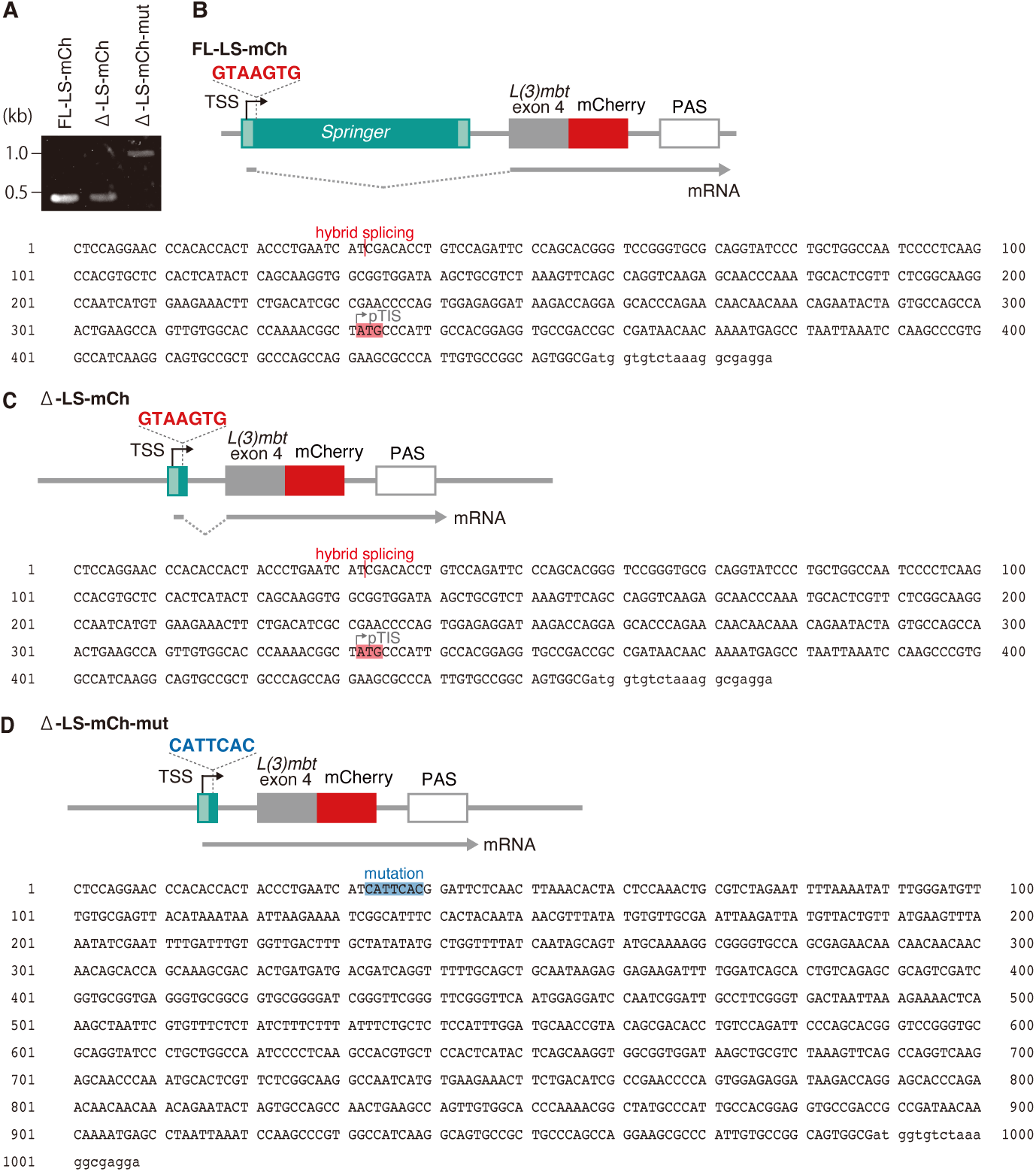
Reporter assays. (A) RT-PCR products of FL-LS-mCh, Δ-LS-mCh, and Δ-LS-mCh-mut mRNAs. (B to D) Sequences of FL-LS-mCh, Δ-LS-mCh, and Δ-LS-mCh-mut mRNAs.

**Fig. S4.**
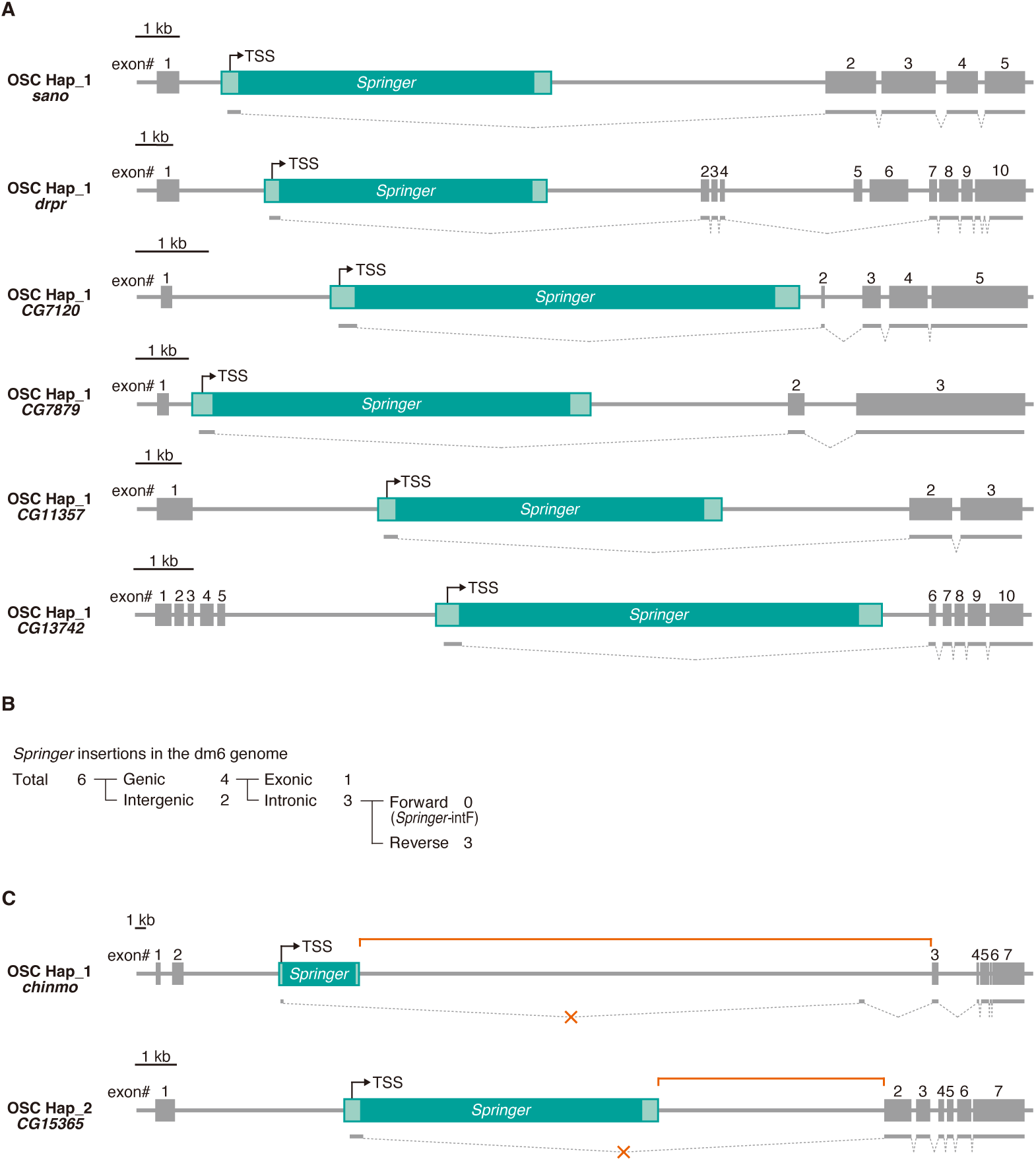
*Springer*-intF insertions within host genes. (A) Six additional examples of *Springer*-intF that drive host-TE hybrid splicing. (B) Genome-wide classification of *Springer* insertions in the dm6 genome. (C) Two additional examples of *Springer*-intF insertions that do not promote hybrid splicing.

**Fig. S5.**
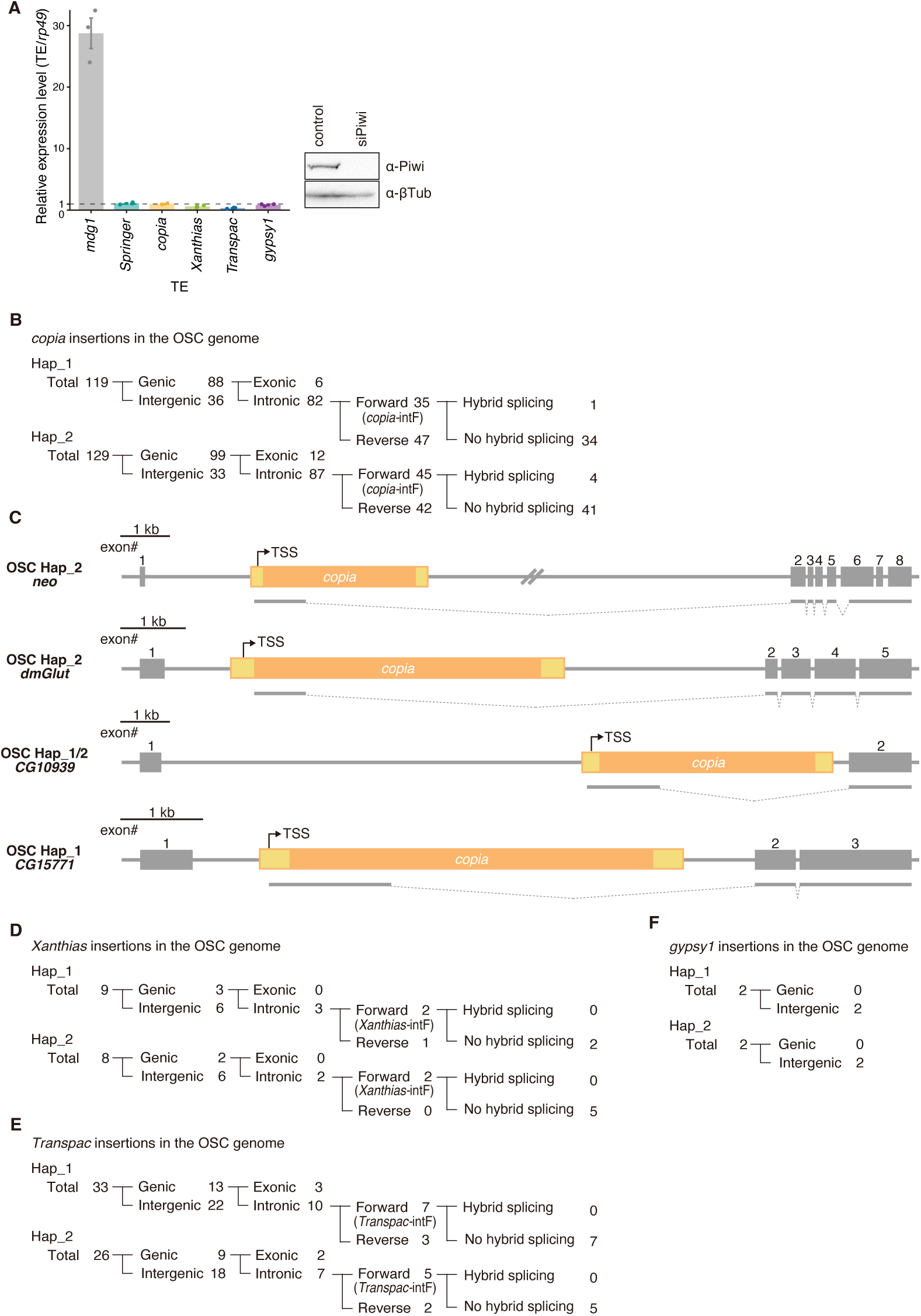
*Copia, Xanthias, Transpac*, and *gypsy1* insertions in OSCs. (A) RT-qPCR showing the expression levels of various TEs in naïve and Piwi-lacking OSCs. Western blot shows the efficiency of Piwi knockdown. (B) Genomic classification of *copia* insertions. (C) *Copia*-intF insertions that promote hybrid splicing in the OSC genome. (D to F) Genomic classification of *Xanthias*, *Transpac*, and *gypsy1* insertions.

**Fig. S6.**
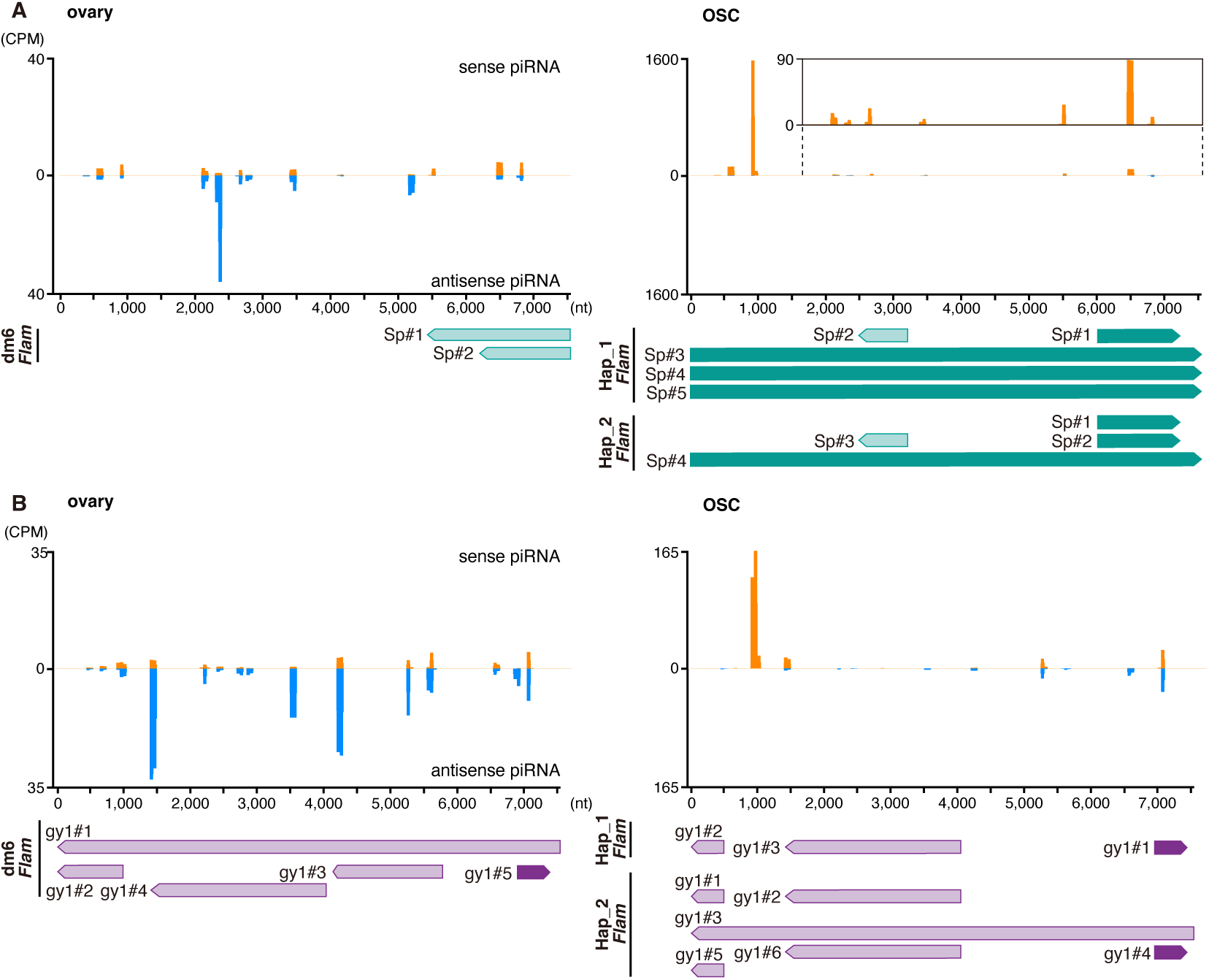
piRNA mapping onto *Springer* and *gypsy1* elements. (A) Mapping of ovarian and OSC *Springer*-piRNAs to the *Springer* consensus. *Springer* fragments annotated in dm6 and within the OSC *Flam* locus are shown below. (B) Mapping of ovarian and OSC *Gypsy1*-piRNAs to the *gypsy1* consensus. *Gypsy1* fragments annotated in dm6 and within the OSC *Flam* locus are shown below.

**Fig. S7.**
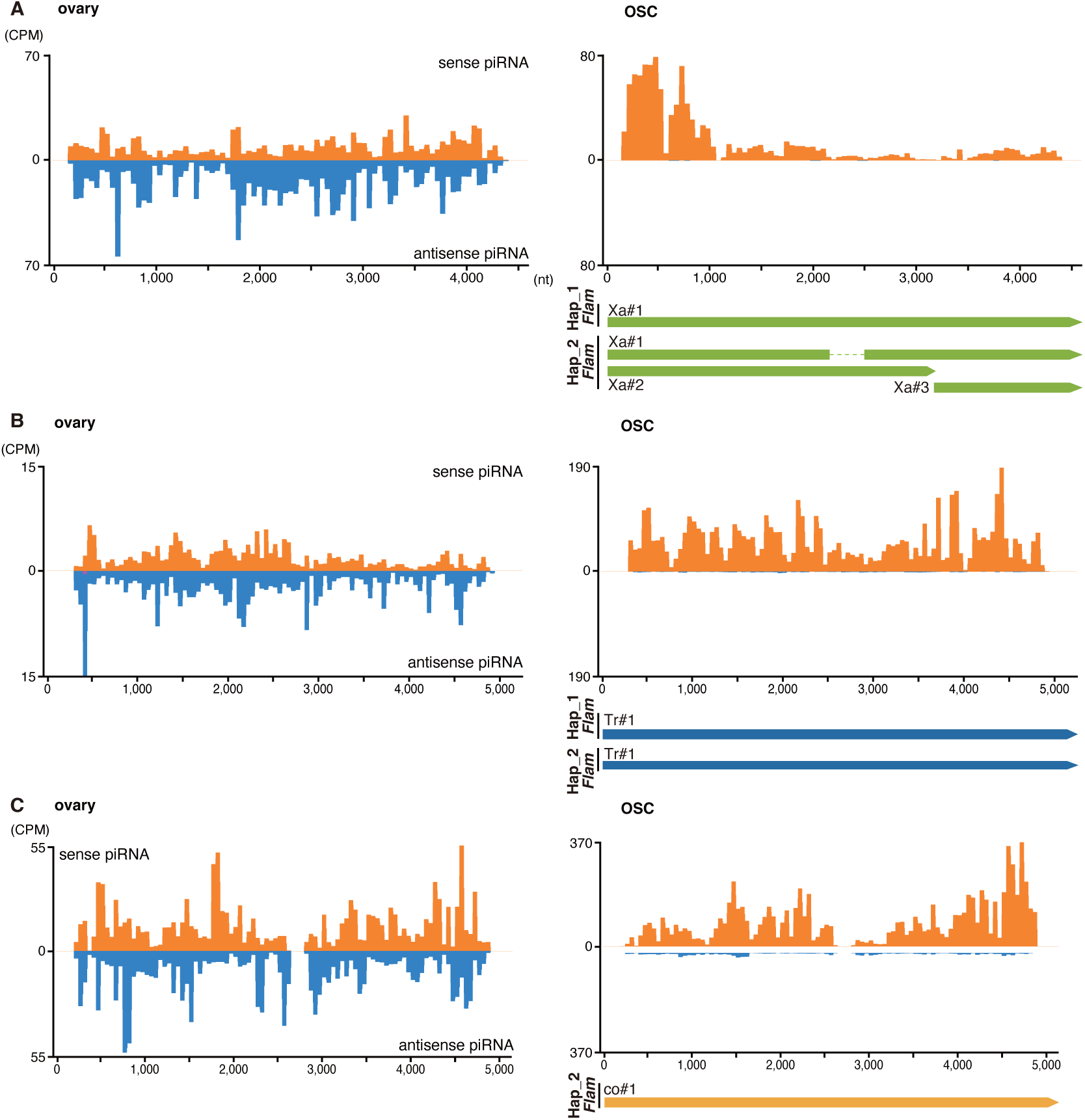
piRNA mapping onto *Xanthias*, *Transpac*, and *copia* elements. (**A**) Mapping of ovarian and OSC *Xanthias*-piRNAs to the *Xanthias* consensus*. Xanthias* fragments annotated in dm6 and within the OSC *Flam* locus are shown below. (**B**) Mapping of ovarian and OSC *Transpac*-piRNAs to the *Transpac* consensus*. Transpac* fragments annotated in dm6 and within the OSC *Flam* locus are shown below. (**C**) Mapping of ovarian and OSC *Copia*-piRNAs to the *copia* consensus*. Copia* fragment annotated in dm6 and within the OSC *Flam* locus is also shown.

**Fig. S8.**
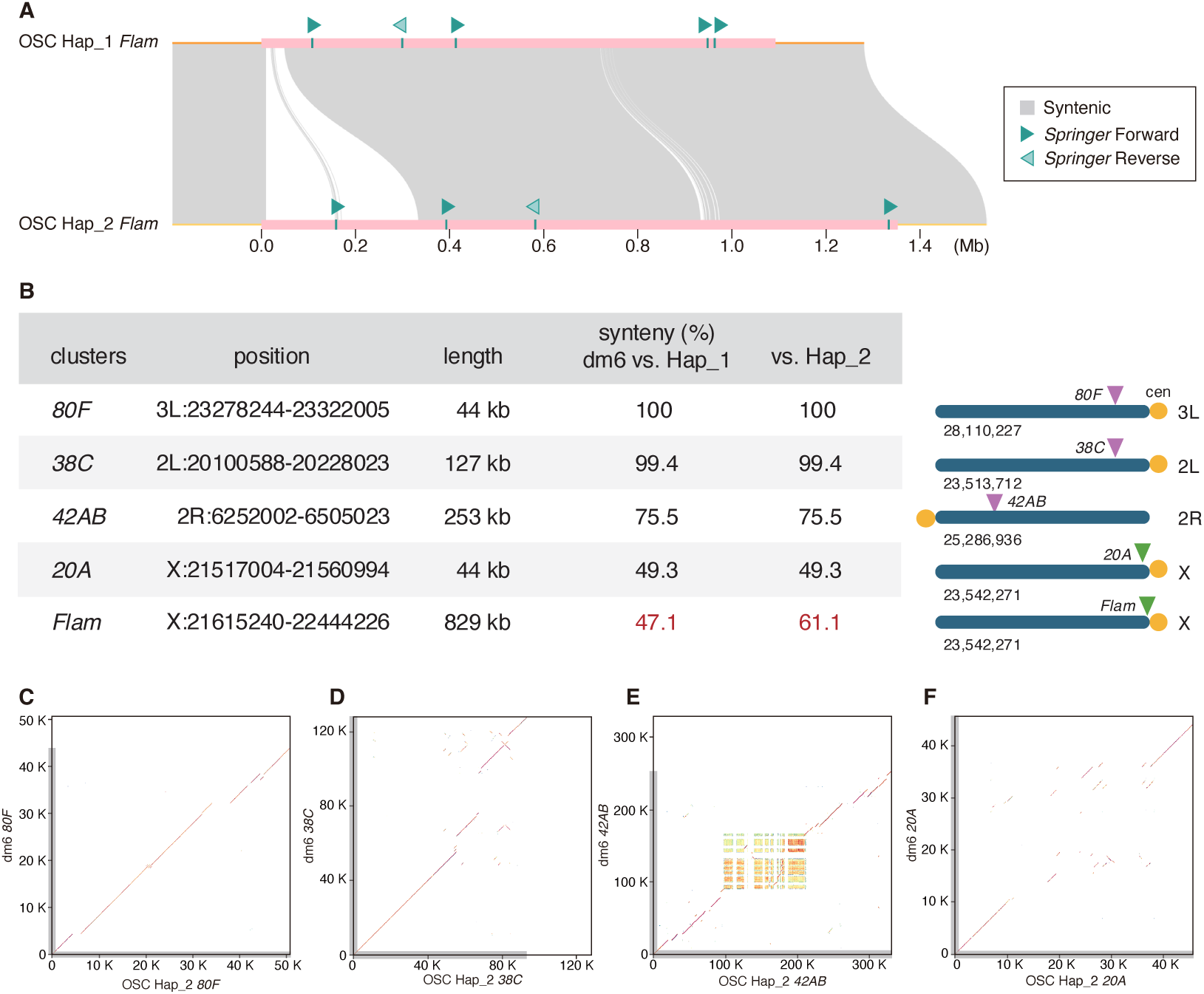
Architecture and comparative genomics of major piRNA clusters. (A) Structural rearrangements within the *Flam* region and its flanking sequences, comparing OSC Hap_1 and Hap_2 assemblies. (B) Summary statistics and annotations for piRNA clusters *80F*, *38C*, *42AB*, *20A*, and *Flam*. (C to F) Pairwise comparisons between dm6 and Hap_2 for *80F* (C), *38C* (D), *42AB* (E), and *20A* (F).

**Fig. S9.**
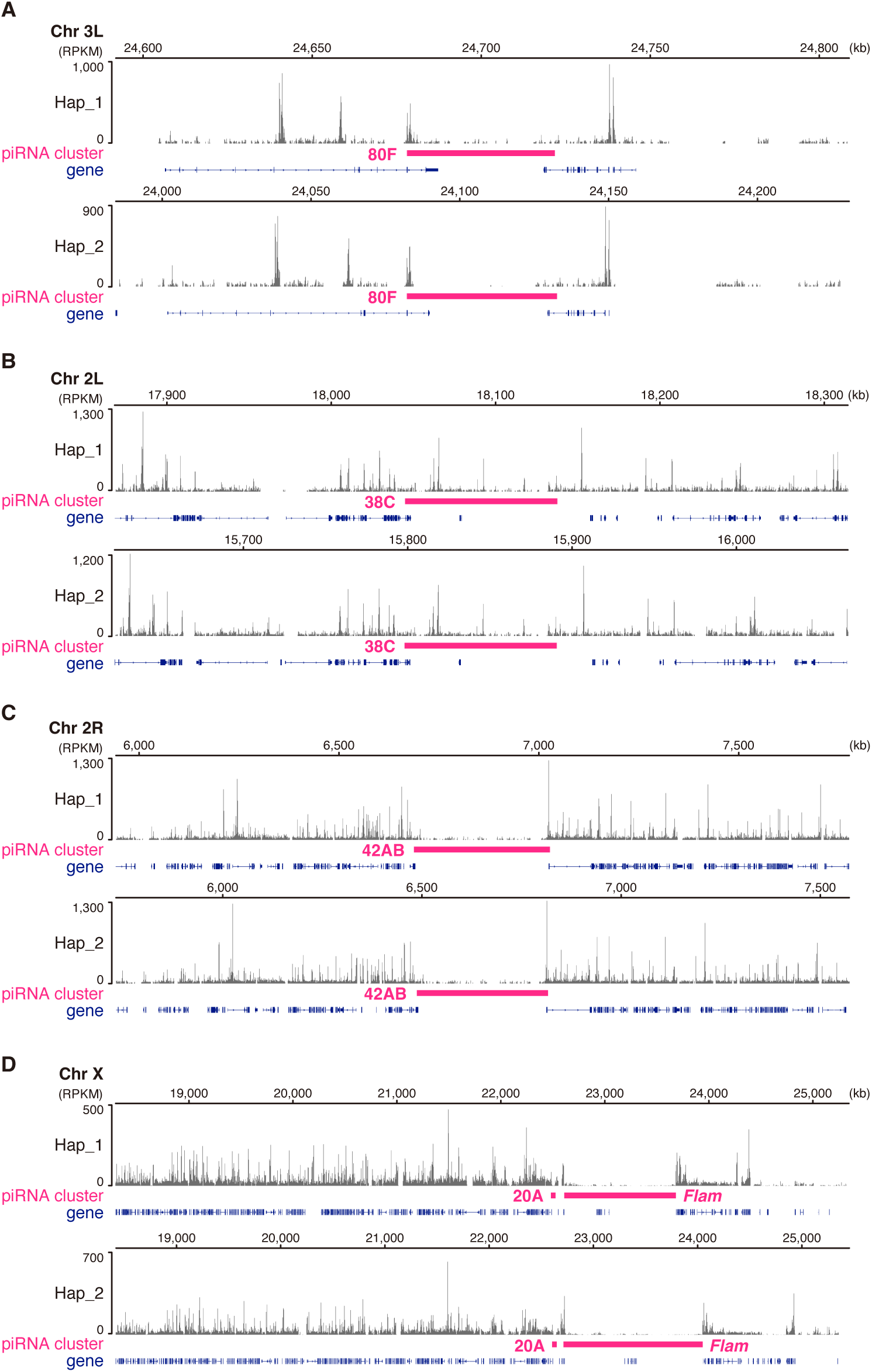
ATAC-seq profiles at major piRNA clusters. (**A** to **D**) ATAC-seq signals across the Hap_1 and Hap_2 assemblies at the indicated clusters: *80F* (A), *38C* (B)*, 42AB* (C), and *20A* (D).

**Fig. S10.**
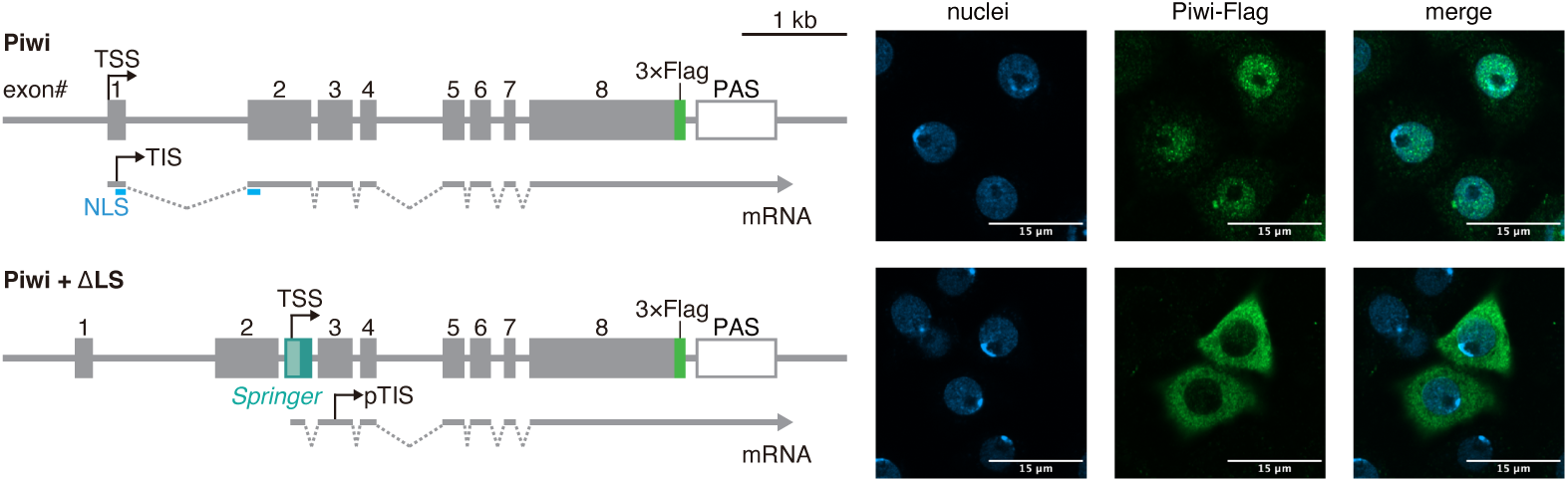
The *Springer* mini cassette converts Piwi to be a cytoplasmic protein. Insertion of the *Springer* mini cassette (Fig. 1D) into *Piwi* intron 2 converts *Piwi* (green) into a cytoplasmic protein. Anti-Flag is used. Blue: DAPI (nuclei).

**Table S1.** *Springer* insertions in the dm6 genome.

| chromosome | start | end | strand | genic/intergenic | exonic/intergenic | sense/anti-sense | host gene | potential to hybrid splicing |
| --- | --- | --- | --- | --- | --- | --- | --- | --- |
| 2L | 3251549 | 3259058 | + | intergenic | . | . | . | × |
| 3L | 20476808 | 20484350 | + | genic | exonic | sense | isoQC | × |
| 3L | 27224890 | 27232435 | + | intergenic | . | . | . | × |
| 3R | 31074284 | 31081826 | - | genic | intronic | anti-sense | Sox100B | × |
| X | 5096328 | 5103837 | - | genic | intronic | anti-sense | CG15465 | × |
| X | 10348854 | 10356363 | + | genic | intronic | anti-sense | I(1)G0289 | × |

**Table S2.** *Springer*-intF elements.

| OSC Hap_1 |  |  |  |  |  |  |  |
| --- | --- | --- | --- | --- | --- | --- | --- |
| chromosome | start | end | strand | genic/intergenic | exonic/intronic | sense/antisense | hybrid splicing |
| chr2L_RagTag | 1760671 | 1768213 | + | genic | intronic | sense | . |
| chr2L_RagTag | 3493199 | 3500742 | + | genic | both intronic | mixed | Ccdc85 |
| chr2L_RagTag | 4079265 | 4086807 | - | genic | intronic | sense | . |
| chr2L_RagTag | 4797512 | 4805055 | + | genic | intronic | sense | . |
| chr2L_RagTag | 5672025 | 5679568 | - | genic | intronic | sense | hoe1 |
| chr2L_RagTag | 5766198 | 5773741 | - | genic | intronic | sense | Rtnl1 |
| chr2L_RagTag | 5855254 | 5862796 | - | genic | intronic | sense | Pgant5 |
| chr2L_RagTag | 10187708 | 10195250 | - | genic | intronic | sense | bun |
| chr2L_RagTag | 12995228 | 13002740 | - | genic | intronic | sense | . |
| chr2L_RagTag | 19184658 | 19192201 | + | genic | intronic | sense | Hr39 |
| chr2R_RagTag | 9680846 | 9688389 | - | genic | intronic | sense | CG13742 |
| chr2R_RagTag | 11673038 | 11680580 | - | genic | intronic | sense | luna |
| chr2R_RagTag | 13095870 | 13103413 | + | genic | both intronic | mixed | . |
| chr2R_RagTag | 14000956 | 14008499 | - | genic | intronic | sense | . |
| chr2R_RagTag | 14810429 | 14817971 | - | genic | intronic | sense | . |
| chr2R_RagTag | 16947018 | 16954561 | + | genic | intronic | sense | lbk |
| chr2R_RagTag | 17525453 | 17532996 | - | genic | intronic | sense | Pgant9 |
| chr2R_RagTag | 19791005 | 19798547 | + | genic | intronic | sense | sano |
| chr2R_RagTag | 25635515 | 25643057 | - | genic | both intronic | mixed | . |
| chr3L_RagTag | 268847 | 276390 | + | genic | intronic | sense | RhoGEF3 |
| chr3L_RagTag | 1568194 | 1575737 | - | genic | intronic | sense | CG7879 |
| chr3L_RagTag | 1736062 | 1743605 | - | genic | intronic | sense | drpr |
| chr3L_RagTag | 2318444 | 2325987 | - | genic | intronic | sense | . |
| chr3L_RagTag | 2845099 | 2852642 | - | genic | both intronic | both sense | . |
| chr3L_RagTag | 4412845 | 4420357 | - | genic | intronic | sense | CG32264 |
| chr3L_RagTag | 5255253 | 5262765 | + | genic | intronic | sense | CG11357 |
| chr3L_RagTag | 5332002 | 5339545 | - | genic | intronic | sense | . |
| chr3L_RagTag | 6090867 | 6098410 | - | genic | both intronic | sense | lama/CG46456 |
| chr3L_RagTag | 6696632 | 6704175 | - | genic | intronic | sense | Rcc1 |
| chr3L_RagTag | 9178326 | 9185839 | + | genic | intronic | sense | CG7120 |
| chr3L_RagTag | 28207762 | 28215305 | + | genic | intronic | sense | . |
| chr3R_RagTag | 2459409 | 2466952 | + | genic | intronic | sense | . |
| chr3R_RagTag | 2505653 | 2513195 | + | genic | intronic | sense | . |
| chr3R_RagTag | 4754275 | 4761818 | + | genic | intronic | sense | CG42402 |
| chr3R_RagTag | 6507995 | 6515537 | - | genic | intronic | sense | CG2082 |
| chr3R_RagTag | 7785641 | 7793182 | - | genic | intronic | sense | Antp |
| chr3R_RagTag | 11829201 | 11836744 | - | genic | intronic | sense | CG14696 |
| chr3R_RagTag | 15371011 | 15378553 | - | genic | intronic | sense | . |
| chr3R_RagTag | 18594317 | 18601859 | - | genic | intronic | sense | Dscam3 |
| chr3R_RagTag | 19587991 | 19595505 | - | genic | intronic | sense | . |
| chr3R_RagTag | 22600237 | 22607780 | + | genic | intronic | sense | . |
| chr3R_RagTag | 22735758 | 22743301 | - | genic | both intronic | mixed | RhoGAP93B |
| chr3R_RagTag | 23017825 | 23025368 | - | genic | intronic | sense | mod(mdg4) |
| chr3R_RagTag | 24498118 | 24505661 | + | genic | mixed | sense | . |
| chr3R_RagTag | 26347030 | 26354572 | - | genic | intronic | sense | REPTOR |
| chr3R_RagTag | 31280289 | 31287831 | + | genic | intronic | sense | Ptp99A |
| chr4_RagTag | 469624 | 477166 | - | genic | intronic | sense | . |
| chrX_RagTag | 1514702 | 1522246 | - | genic | intronic | sense | . |
| chrX_RagTag | 2599876 | 2607418 | + | genic | intronic | sense | . |
| chrX_RagTag | 4014246 | 4021789 | + | genic | intronic | sense | ec |
| chrX_RagTag | 4278495 | 4286038 | + | genic | intronic | sense | . |
| chrX_RagTag | 8521396 | 8528939 | + | genic | intronic | sense | sdt |
| chrX_RagTag | 10041112 | 10048655 | - | genic | intronic | sense | Ptpmeg2 |
| chrX_RagTag | 11286948 | 11294491 | - | genic | intronic | sense | . |
| chrX_RagTag | 12735462 | 12742975 | + | genic | intronic | sense | . |
| chrX_RagTag | 13934407 | 13941949 | + | genic | intronic | sense | . |
| chrX_RagTag | 17439484 | 17447026 | - | genic | intronic | sense | goe |
| chrX_RagTag | 19957298 | 19964840 | - | genic | intronic | sense | RhoGAP18B |
| chrX_RagTag | 20109758 | 20117300 | - | genic | intronic | sense | CG8034 |
| chrX_RagTag | 20591714 | 20599257 | - | genic | intronic | sense | lncRNA:CR44885 |
| chrX_RagTag | 21963392 | 21970935 | + | genic | intronic | sense | bves |
| chrX_RagTag | 21972771 | 21980314 | + | genic | intronic | sense | . |
| chrX_RagTag | 21982205 | 21989748 | + | genic | intronic | sense | bves |

| OSC Hap_2 |  |  |  |  |  |  |  |
| --- | --- | --- | --- | --- | --- | --- | --- |
| chromosome | start | end | strand | genic/intergenic | exonic/intronic | sense/antisense | hybrid splicing |
| chr2L_RagTag | 1741661 | 1749204 | + | genic | intronic | sense | . |
| chr2L_RagTag | 3920631 | 3928145 | + | genic | intronic | sense | . |
| chr2L_RagTag | 3928154 | 3935668 | + | genic | intronic | sense | . |
| chr2L_RagTag | 4728208 | 4735751 | - | genic | intronic | sense | Cf2 |
| chr2L_RagTag | 4786700 | 4794243 | - | genic | intronic | sense | hoe1 |
| chr2L_RagTag | 7382965 | 7390508 | - | genic | intronic | sense | . |
| chr2L_RagTag | 7698520 | 7706062 | - | genic | intronic | sense | Samuel |
| chr2L_RagTag | 8958373 | 8965916 | + | genic | intronic | sense | bru1 |
| chr2L_RagTag | 14135346 | 14142879 | - | genic | intronic | sense | . |
| chr2L_RagTag | 16059860 | 16067403 | - | genic | intronic | sense | CG10947 |
| chr2R_RagTag | 1317846 | 1325389 | + | genic | intronic | sense | . |
| chr2R_RagTag | 6338498 | 6346041 | - | genic | intronic | sense | EcR |
| chr2R_RagTag | 6355828 | 6363371 | - | genic | intronic | sense | EcR |
| chr2R_RagTag | 7289601 | 7297142 | - | genic | intronic | sense | mim |
| chr2R_RagTag | 11568116 | 11575658 | - | genic | intronic | sense | luna |
| chr2R_RagTag | 15114879 | 15122422 | + | genic | intronic | sense | PRAS40 |
| chr2R_RagTag | 16658925 | 16666467 | - | genic | intronic | sense | Zasp52 |
| chr2R_RagTag | 17081037 | 17088580 | + | genic | intronic | sense | . |
| chr2R_RagTag | 25737652 | 25745194 | - | genic | both intronic | mixed | CG4622 |
| chr3L_RagTag | 1431325 | 1438867 | + | genic | intronic | sense | . |
| chr3L_RagTag | 1444595 | 1452138 | + | genic | intronic | sense | ru |
| chr3L_RagTag | 4057015 | 4064527 | - | genic | intronic | sense | CG32264 |
| chr3L_RagTag | 5716850 | 5724393 | - | genic | intronic | sense | lama |
| chr3L_RagTag | 8487859 | 8495402 | + | genic | intronic | sense | nmo |
| chr3L_RagTag | 8872875 | 8880388 | + | genic | intronic | sense | CG7120 |
| chr3L_RagTag | 9349452 | 9356994 | + | genic | intronic | sense | dally |
| chr3L_RagTag | 11193855 | 11201398 | + | genic | intronic | sense | simj |
| chr3L_RagTag | 15338629 | 15346171 | + | genic | intronic | sense | ome |
| chr3L_RagTag | 16214564 | 16222107 | + | genic | intronic | sense | CrebA |
| chr3L_RagTag | 20027328 | 20034871 | - | genic | intronic | sense | fz2 |
| chr3L_RagTag | 31037647 | 31045190 | - | genic | intronic | sense | . |
| chr3R_RagTag | 11020915 | 11028428 | - | genic | intronic | sense | CG9837 |
| chr3R_RagTag | 11135357 | 11142900 | - | genic | intronic | sense | . |
| chr3R_RagTag | 11683411 | 11690954 | + | genic | intronic | sense | . |
| chr3R_RagTag | 19900731 | 19908274 | - | genic | intronic | sense | cher |
| chr3R_RagTag | 21244947 | 21252459 | - | genic | intronic | sense | CG6231 |
| chr3R_RagTag | 23461942 | 23469485 | + | genic | intronic | sense | . |
| chr3R_RagTag | 27837865 | 27845408 | + | genic | intronic | sense | . |
| chr3R_RagTag | 28169969 | 28177482 | - | genic | intronic | sense | l(3)mbt |
| chr3R_RagTag | 30760826 | 30768369 | + | genic | intronic | sense | . |
| chr3R_RagTag | 30850489 | 30858032 | - | genic | intronic | sense | Atg16 |
| chr3R_RagTag | 31964865 | 31972408 | - | genic | intronic | sense | Sap-r |
| chr3R_RagTag | 34168037 | 34175580 | - | genic | intronic | sense | Akap200 |
| chr4_RagTag | 565219 | 572761 | - | genic | intronic | sense | . |
| chrX_RagTag | 1404194 | 1411738 | - | genic | intronic | sense | . |
| chrX_RagTag | 2483159 | 2490701 | + | genic | intronic | sense | Raf |
| chrX_RagTag | 4947310 | 4954853 | + | genic | intronic | sense | CG32772 |
| chrX_RagTag | 5402846 | 5410389 | + | genic | intronic | sense | SK |
| chrX_RagTag | 6011632 | 6019175 | + | genic | intronic | sense | Act5C |
| chrX_RagTag | 6054614 | 6062157 | + | genic | intronic | sense | . |
| chrX_RagTag | 8380252 | 8387794 | + | genic | intronic | sense | sdt |
| chrX_RagTag | 9282953 | 9290496 | - | genic | intronic | sense | . |
| chrX_RagTag | 11279788 | 11287331 | - | genic | intronic | sense | . |
| chrX_RagTag | 11316047 | 11323590 | - | genic | intronic | sense | . |
| chrX_RagTag | 12715119 | 12722662 | + | genic | intronic | sense | . |
| chrX_RagTag | 17491814 | 17506520 | - | genic | intronic | sense | . |
| chrX_RagTag | 19679300 | 19686842 | + | genic | intronic | sense | S6KL |

**Table S3.**
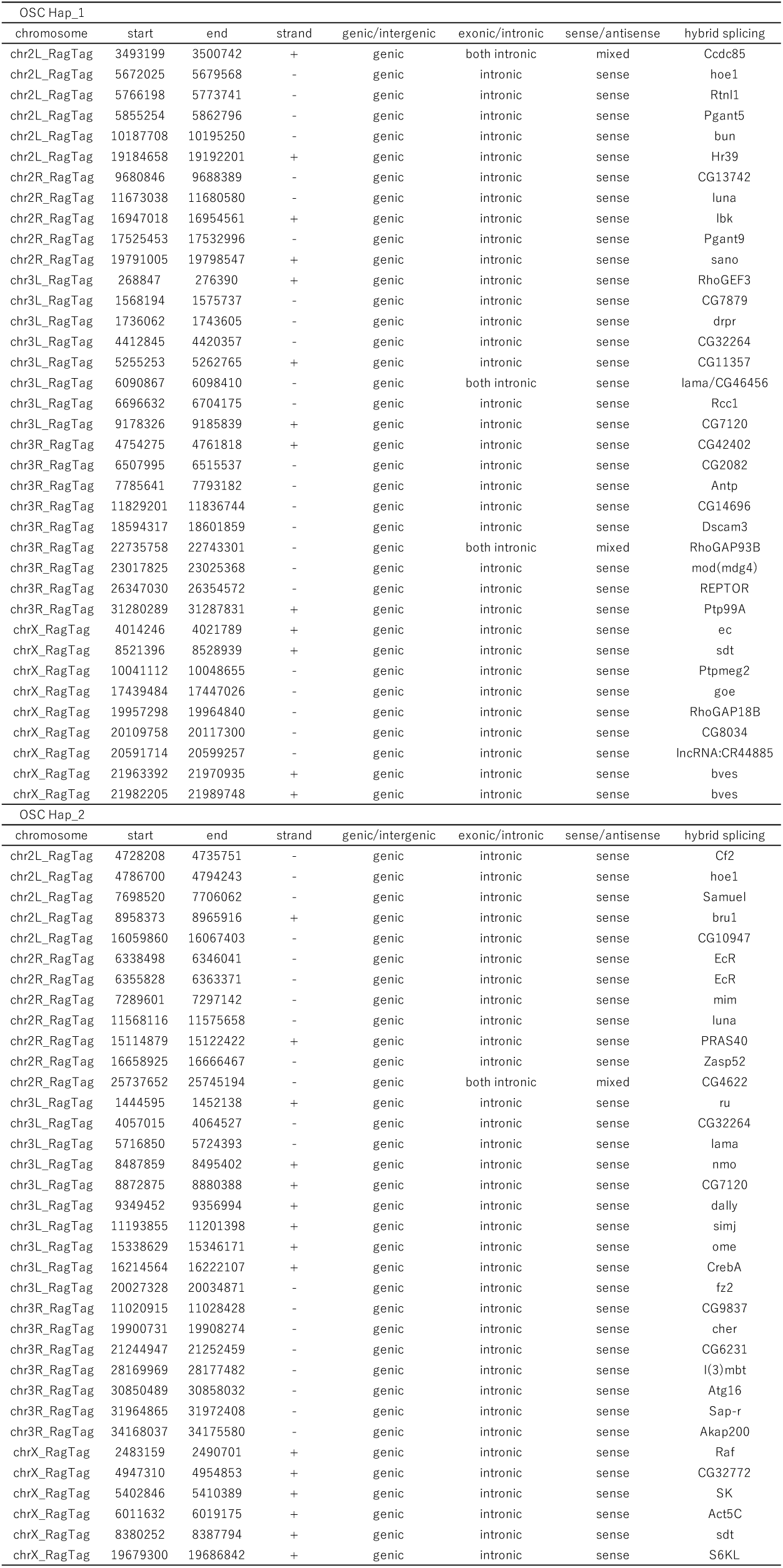
*Springer*-intF *elements that promote hybrid splicing*.

**Table S4.** *Springer*-intF *elements that do not promote hybrid splicing*.

| OSC Hap_1 |  |  |  |  |  |  |
| --- | --- | --- | --- | --- | --- | --- |
| chromosome | start | end | strand | genic/intergenic | exonic/intronic | sense/antisense |
| chr2L_RagTag | 1760671 | 1768213 | + | genic | intronic | sense |
| chr2L_RagTag | 4079265 | 4086807 | - | genic | intronic | sense |
| chr2L_RagTag | 4797512 | 4805055 | + | genic | intronic | sense |
| chr2L_RagTag | 12995228 | 13002740 | - | genic | intronic | sense |
| chr2R_RagTag | 13095870 | 13103413 | + | genic | both intronic | mixed |
| chr2R_RagTag | 14000956 | 14008499 | - | genic | intronic | sense |
| chr2R_RagTag | 14810429 | 14817971 | - | genic | intronic | sense |
| chr2R_RagTag | 25635515 | 25643057 | - | genic | both intronic | mixed |
| chr3L_RagTag | 2318444 | 2325987 | - | genic | intronic | sense |
| chr3L_RagTag | 2845099 | 2852642 | - | genic | both intronic | both sense |
| chr3L_RagTag | 5332002 | 5339545 | - | genic | intronic | sense |
| chr3L_RagTag | 28207762 | 28215305 | + | genic | intronic | sense |
| chr3R_RagTag | 2459409 | 2466952 | + | genic | intronic | sense |
| chr3R_RagTag | 2505653 | 2513195 | + | genic | intronic | sense |
| chr3R_RagTag | 15371011 | 15378553 | - | genic | intronic | sense |
| chr3R_RagTag | 19587991 | 19595505 | - | genic | intronic | sense |
| chr3R_RagTag | 22600237 | 22607780 | + | genic | intronic | sense |
| chr3R_RagTag | 24498118 | 24505661 | + | genic | mixed | sense |
| chr4_RagTag | 469624 | 477166 | - | genic | intronic | sense |
| chrX_RagTag | 1514702 | 1522246 | - | genic | intronic | sense |
| chrX_RagTag | 2599876 | 2607418 | + | genic | intronic | sense |
| chrX_RagTag | 4278495 | 4286038 | + | genic | intronic | sense |
| chrX_RagTag | 11286948 | 11294491 | - | genic | intronic | sense |
| chrX_RagTag | 12735462 | 12742975 | + | genic | intronic | sense |
| chrX_RagTag | 13934407 | 13941949 | + | genic | intronic | sense |
| chrX_RagTag | 21972771 | 21980314 | + | genic | intronic | sense |
| OSC Hap_2 |  |  |  |  |  |  |
| chromosome | start | end | strand | genic/intergenic | exonic/intronic | sense/antisense |
| chr2L_RagTag | 1741661 | 1749204 | + | genic | intronic | sense |
| chr2L_RagTag | 3920631 | 3928145 | + | genic | intronic | sense |
| chr2L_RagTag | 3928154 | 3935668 | + | genic | intronic | sense |
| chr2L_RagTag | 7382965 | 7390508 | - | genic | intronic | sense |
| chr2L_RagTag | 14135346 | 14142879 | - | genic | intronic | sense |
| chr2R_RagTag | 1317846 | 1325389 | + | genic | intronic | sense |
| chr2R_RagTag | 17081037 | 17088580 | + | genic | intronic | sense |
| chr3L_RagTag | 1431325 | 1438867 | + | genic | intronic | sense |
| chr3L_RagTag | 31037647 | 31045190 | - | genic | intronic | sense |
| chr3R_RagTag | 11135357 | 11142900 | - | genic | intronic | sense |
| chr3R_RagTag | 11683411 | 11690954 | + | genic | intronic | sense |
| chr3R_RagTag | 23461942 | 23469485 | + | genic | intronic | sense |
| chr3R_RagTag | 27837865 | 27845408 | + | genic | intronic | sense |
| chr3R_RagTag | 30760826 | 30768369 | + | genic | intronic | sense |
| chr4_RagTag | 565219 | 572761 | - | genic | intronic | sense |
| chrX_RagTag | 1404194 | 1411738 | - | genic | intronic | sense |
| chrX_RagTag | 6054614 | 6062157 | + | genic | intronic | sense |
| chrX_RagTag | 9282953 | 9290496 | - | genic | intronic | sense |
| chrX_RagTag | 11279788 | 11287331 | - | genic | intronic | sense |
| chrX_RagTag | 11316047 | 11323590 | - | genic | intronic | sense |
| chrX_RagTag | 12715119 | 12722662 | + | genic | intronic | sense |
| chrX_RagTag | 17491814 | 17506520 | - | genic | intronic | sense |

**Table S5.** Other clusters that produce *Transpac-*, *Xanthias-*, and *copia*-piRNAs in ovaries.

| copia |  |  |  |  |  |  |  |  |
| --- | --- | --- | --- | --- | --- | --- | --- | --- |
| chromosome | start | end | TE name | strand | chromosome | start | end | piRNA cluster name |
| 2L | 20155808 | 20156133 | copia | + | 2L | 20100587 | 20228023 | 38C |
| 2L | 20157412 | 20157737 | copia | + | 2L | 20100587 | 20228023 | 38C |
| 2L | 20159016 | 20164268 | copia | + | 2L | 20100587 | 20228023 | 38C |
| 2R | 6449296 | 6450137 | copia | + | 2R | 6252001 | 6505023 | 42AB |
| 2R | 6450185 | 6450672 | copia | + | 2R | 6252001 | 6505023 | 42AB |
| 3L | 24307624 | 24308621 | copia | - | 3L | 24302050 | 24329391 | . |
| 3L | 24308912 | 24309792 | copia | - | 3L | 24302050 | 24329391 | . |
| 3L | 24513400 | 24514280 | copia | - | 3L | 24491315 | 24519769 | . |
| 3R | 4151653 | 4151988 | copia | + | 3R | 4126770 | 4152021 | . |
| Xanthias |  |  |  |  |  |  |  |  |
| chromosome | start | end | TE name | strand | chromosome | start | end | cluster name |
| 2R | 6398093 | 6402626 | Xanthias | + | 2R | 6252001 | 6505023 | 42AB |
| 3L | 25611531 | 25612125 | Xanthias | - | 3L | 25607998 | 25617999 | . |
| 3L | 25613300 | 25614005 | Xanthias | - | 3L | 25607998 | 25617999 | . |
| X | 21528329 | 21528680 | Xanthias | - | X | 21517003 | 21560994 | 20A |
| Transpac |  |  |  |  |  |  |  |  |
| chromosome | start | end | TE name | strand | chromosome | start | end | cluster name |
| 3L | 26704581 | 26709829 | Transpac | + | 3L | 26694437 | 26779349 | . |

**Table S6.** Locus information.

|  |  |
| --- | --- |
| Flam |  |
| genome | locus |
| dm6 | X:21615240-22444226 |
| OSC Hap_1 | chrX_RagTag:22608911-23685678 |
| OSC Hap_2 | chrX_RagTag:22712449-24050631 |
| 20A |  |
| genome | locus |
| dm6 | X:21517004-21560994 |
| OSC Hap_1 | chrX_RagTag:22483951-22529500 |
| OSC Hap_2 | chrX_RagTag:22604372-22649920 |
| 38C |  |
| genome | locus |
| dm6 | 2L:20100588-20228023 |
| OSC Hap_1 | chr2L_RagTag:18045121-18138016 |
| OSC Hap_2 | chr2L_RagTag:15797893-15890788 |
| 42AB |  |
| genome | locus |
| dm6 | 2R:6252002-6505023 |
| OSC Hap_1 | chr2R_RagTag:6689199-7029960 |
| OSC Hap_2 | chr2R_RagTag:6488019-6816562 |
| 80F |  |
| genome | locus |
| dm6 | 3L:23278244-23322005 |
| OSC Hap_1 | chr3L_RagTag:24678298-24722059 |
| OSC Hap_2 | chr3L_RagTag:24082398-24132906 |

**Table S7.**
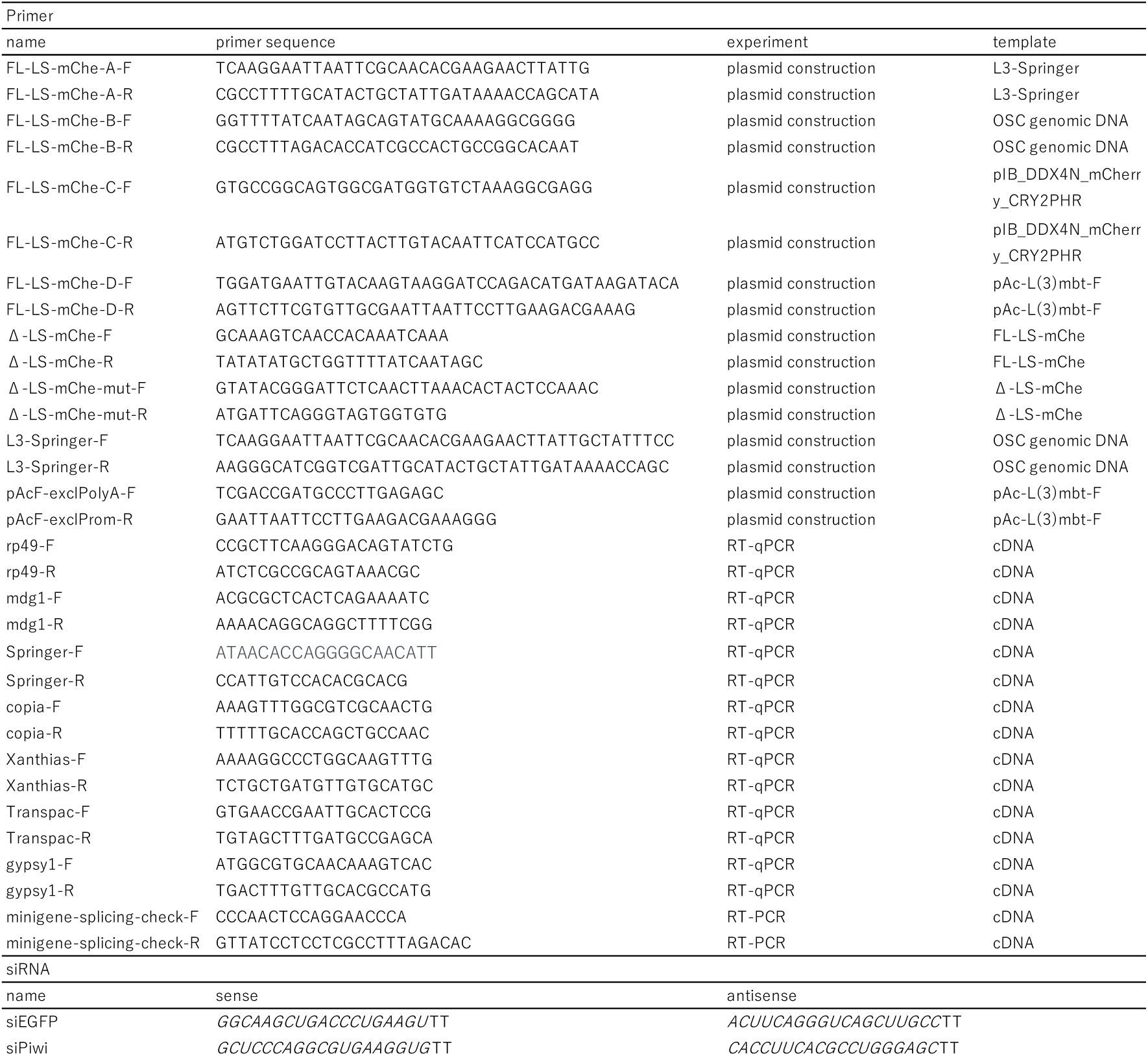
Primer and siRNA sequences.

